# Spatiotemporal Dynamics and Environmental Drivers of Maize Microbiome Structure across Diverse Agroecosystems in Georgia, USA

**DOI:** 10.64898/2026.11.19.767978

**Authors:** Aduragbemi Amo, Chen Yang, Matthew S. Johnson, Lynsey Kovar, Jason G. Wallace

**Author notes:** Current address: Seattle, Washington, USA. Current address: College Station, Texas, USA.

## Abstract

The understanding of how microbial communities differ across soils, environments, plant tissues, and developmental stages is key to the improvement of crop productivity and resilience. Here, we conducted a statewide survey of maize (*Zea mays* L.) microbiomes across 35 fields at 16 locations in Georgia, USA, sampling soil, rhizosphere, root interior, and stalk interior at two developmental stages (600 and 1400 growing degree days, GDD). More than 38000 ASVs were produced after amplicon sequencing of 16S rRNA genes. The phyla shown to be dominant are Proteobacteria, Actinobacteriota, Chloroflexi, Planctomycetota*, and* Acidobacteriota. Based on alpha and beta diversity analyses, the below-ground compartments (soil and rhizosphere) revealed higher diversity and stronger spatial structuring when compared to the above-ground sample types, whereas root and stalk interior microbiomes revealed marked shifts across growth stages. Amino acid metabolism, secondary metabolite biosynthesis, and energy-related pathways predictions (via PICRUSt2) show stage-dependent enrichment. Network analyses show persistent keystone taxa (Proteobacteria and Actinobacteriota) mediating microbial stability, coupled with an increase in modularity and connectivity as we move from early to late growth stages. Some environmental variables, such as temperature, humidity, soil potassium, and soil sodium, showed significant correlation with microbial diversity indicative of a compartment-specific environmental sensitivity. Collectively, our results show that in maize, microbial community composition, interaction, and potential functionality are both impacted by developmental stages and environmental heterogeneity. This study provides a fundamental framework for understanding the microbiome dynamics in field-grown maize across various agroecological conditions. It equally provides ecological insights relevant to microbiome-informed crop management.

## Introduction

Plant microbiomes consist of microorganisms such as bacteria, fungi, algae, protozoa, and viruses, all of which can be beneficial, neutral, or pathogenic toward the plant (Mendes et al., 2013). The microbiomes associated with plants can be divided into several compartments; the most common of these are bulk soil, meaning soil not influenced by roots; the rhizosphere, meaning soil immediately adjacent to roots that is impacted by dead plant cells, exudates from roots, and mucilage (Hartmann et al., 2008); and the endosphere, meaning the inner tissues of a plant, including its leaves, stems, roots, and seeds.

Beneficial microbial communities can enhance the health, growth, and development of plants, including stress tolerance and pathogen resistance (Brader et al., 2017; Hardoim et al., 2015; Lemanceau et al., 2017; Singh & Goodwin, 2022). For example, 18 of the 29 essential elements required by plants are derived from the soil, with soil microbial communities playing a key role in the transfer of these nutrients (Brevik et al., 2020). Soil microbiomes can also promote plant growth hormone synthesis, compete with pathogens for resources, and produce chemicals that induce plant resistance (Ali et al., 2023; Yang et al., 2017a). Rhizosphere microbiomes, meanwhile, can improve seed germination, plant metabolism, nutrient and water uptake, hormonal balance, and stress tolerance (Berendsen et al., 2012; Lugtenberg, 2014; Trivedi et al., 2020). In addition, endophytes can enhance plant growth and provide protection to the plants (Burragoni & Jeon, 2021; Chaudhary et al., 2023; Watts et al., 2023).

Maize (*Zea mays* L.) is a globally important cereal crop, with cereal global production increasing by 61 million tonnes between 2022 and 2023 driven by an increase in maize output (FAO, 2024). An increasing body of research is revealing how maize and its microbiome influence each other, although in most cases the mechanisms are still unclear. For example, soil properties, plant growth stage, and root exudates shape maize’s rhizosphere microbiome (Bai et al., 2022; Brusetti et al., 2005; Dematheis et al., 2012; Emmett et al., 2020; Gomes et al., 2001; Johnston-Monje et al., 2014; Pascale et al., 2020; Tao et al., 2017). Plant genetics also plays a role (Meier et al., 2022; Peiffer et al., 2013; Wallace et al., 2018; Walters et al., 2018), presumably due to genotype-specific differences in plant structure. The maize microbiome also appears to play some role in heterosis (Wagner et al., 2020; Wagner et al., 2021), the phenomenon whereby hybrid maize varieties perform significantly better than inbred parents.

Techniques like microbiome engineering, biofertilizers, and microbial inoculants offers new opportunities to boost maize productivity and sustainability (Khan et al., 2019; Li et al., 2022; Qiu et al., 2019). Despite these advances, research on the maize microbiome is still relatively young (Singh & Goodwin, 2022). Many questions remain about the specific interactions between maize and its microbes, especially since most studies use inbred lines or production practices that do not reflect the actual conditions used for most maize production. To address this issue, we conducted a field survey of maize-associated microbiomes across the state of Georgia. Although Georgia constitutes only small amount of total maize production in the US (0.55%) (U.S. Department of Agriculture (USDA), 2025), it is the 8^th^ most valuable commodity in the state (Kane, 2023), and the variety of soil and climactic conditions across the state allows a wide survey of conditions. Additionally, as climate change progresses, the main corn-growing sections of the US are going to increasingly experience hotter, more stressful growing conditions similar to what corn in Georgia is already experiencing. Our study encompasses soil, rhizosphere, root, and stalk microbiomes collected from field-grown commercial maize at 35 fields at 16 locations across the state of Georgia. Our primary goal was to determine the patterns of microbial diversity associated with maize across the state, with a secondary goal of identifying potential relationships between local environmental conditions and these microbial communities.

## Materials and Methods

Samples were collected from 35 field sites around the state of Georgia during the summer of 2018. Permission was obtained from growers, along with key agronomic information like planting time. Sampling times were targeted based on cumulative growing-degree days (GDDs) to coincide with key windows at the beginning of ear development (V7-12, ∼600 GDDs) and when kernel size was being determined (R1-2, ∼1400 GDDs; Mark Westgate, personal communication). For logistical reasons, samplings were grouped based on which locations were near each other and at approximately the same development. At each location and timepoint, samples were collected of the following types (**Figure 1**): *Between-row soil:* 10 soil cores taken from between corn rows to a depth of appx. 15 cm, homogenized and stored on ice. The soil corer interior was wiped down with 70% isopropyl alcohol between samplings to remove carryover. *In-row soil:* Same as between-row soil, but taken in line with the corn plants. *Rhizosphere/Roots:* Two maize plants 2-4 meters apart were selected and dug up with a drain spade. The roots were vigorously shaken to remove loose soil, then an 8-10 cm segment of root was cut from the root ball with a fresh razor blade and placed in a labeled, sterile container on dry ice. *Stalk:* On the same 2 plants, the stalk was measured and an 8-cm segment cut from its middle (as measured from ground to top). Cut segments were then placed in a clean, labeled plastic bag on dry ice for transport back to the lab. Clippers were wiped down with 70% isopropyl alcohol between samplings to remove any potential carryover. All samples were stored on dry ice for transport back to the lab, then stored at -80°C until processed. GPS coordinates of each sample were recorded at the first (GDD 600) sample time so that the second (GDD 1400) sampling would be nearby (<10 meters apart).

**Figure 1:**
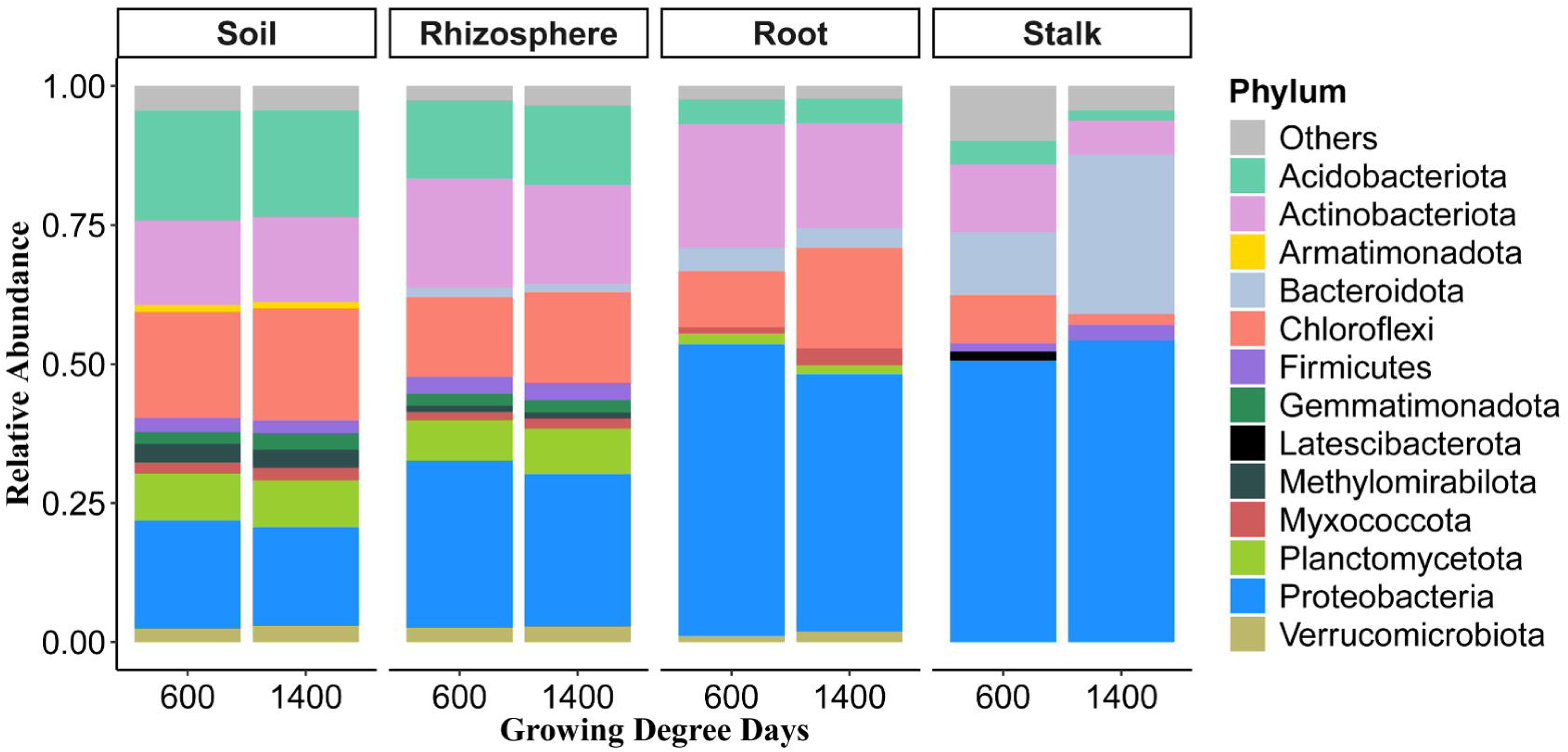

### Soil Analysis

After removing a small amount of homogenized soil for DNA extraction (below), the remainder of each soil sample was sent to the UGA Agricultural and Environmental Services Laboratory for standard analysis.

### Sample Processing

Each sample type was extracted as indicated below. All surfaces and tools were cleaned with 70% ethanol between samples to remove carryover. Blank samples were made by performing the same procedure in the absence of actual sample.

### Soil

Initial, homogenized soil samples were thawed on ice and a pea-sized subsample transferred to a BashingBean Lysis tube (Zymo Quick-DNA Fecal/Soil Microbe 96 kit), then stored at -80°C until extraction.

### Roots & Rhizosphere

Frozen roots (with attached rhizosphere soil) were retrieved from -80° C storage and suspended in 10 mL of deionized water. Rhizosphere soil was collected by vortexing for 15 seconds, after which roots were transferred to new 50 mL tubes using sterilized forceps. The rhizosphere suspension was centrifuged at 4,500xg for 12 minutes, the supernatant then removed via serological pipette, and the remaining pellet stored at -80°C until extraction.

The roots were washed two additional times in 10 mL deionized water to remove any remaining soil, then diced into 1–3 mm fragments using sterile razor blades. Roughly 0.5 mL of root material was then transferred to a 2 mL microcentrifuge tube containing a stainless steel bead, placed immediately on dry ice, then stored at -80° C. After the initial tubes were found to crack during sample grinding (below), any affected samples were re-processed from residual material, and all other root samples were thawed for 15 minutes, transferred to a fresh grind-compatible tube, and refrozen.

### Stalks

Stalk samples were kept at -80° C until processing and maintained on dry ice during handling. For each sample, 2 cm from the end were cut off using a fresh razor blade and discarded, and the next 2-3 cm cut off for sampling. The outer (cortex) layers were removed, and the remaining pith was diced into 1–3 mm fragments. Approximately 1 mL of the processed tissue was transferred into a sterile 2 mL microcentrifuge tube containing a stainless steel bead, which was stored at -80° C. When the initial tubes were found to crack during grinding (below), samples were briefly thawed for 15 minutes, transferred to a fresh (and more robust) sample tube, and refrozen.

### DNA Extraction

All samples were extracted with a Zymo Quick-DNA Fecal/Soil Microbe 96 kit, with slight protocol modifications. 500 μL of BashingBead buffer was added to each tube, which were then ground in a Geno/Grinder at 1400 RMP for 5 minutes. Samples were centrifuged for 8 minutes at 10,000xg to pellet debris, and 250 μL of supernatant transferred to the 96-well block. Extraction then proceeded according to the manufacturer’s protocol, save that centrifuge speeds were set at 2576xg (the maximum for the available equipment), and in the last step the eluent was placed back on the Silicon-A binding plate and centrifuged a second time. For one plate, lysis debris clogged the filters in the initial (Silicon-A plate) binding step; in these cases, a fresh, sterile pipette tip was manually used to scrape debris from the filter to allow the lysate to enter.

### Library Preparation and Sequencing

Library preparation consisted of two sequential PCR amplifications. The first amplification amplified the 16s rRNA gene and added Illumina barcode linkers for the second step. 16s rRNA gene amplification primers were based off Earth Microbiome Project universal primers 515F (Parada et al., 2016) and 806R (Apprill et al., 2015) with staggered frameshift spacers (Lundberg et al., 2013). Final primer sequences were [TCGTCGGCAGCGTCAGATGTGTATAAGAGACAG][X XX][GA][GTGYCAGCMGCCGCGGTAA] (forward) and [GTCTCGTGGGCTCGGAGATGT GTATAAGAGACAG][XXX][GA][GGACTACNVGGGTWTCTAAT] (reverse); bracketed segments indicate (in order) Illumina index adaptors, a 0-5 bp frameshift (denoted XXX and consisting of no insert, T, CT, ACT, GACT, or TGACT), a 2-bp GA adaptor, and the actual 16s amplification primers. Primers were mixed into separate Forward and Reverse primer master mixes, each consisting of the 6 possible forward or reverse primers mixed equimolarly (0.5 μM per primer, or 3 μM for the entire master mix). Organellar sequences were excluded by including peptide nucleic acid blockers (PNA Bio) against chloroplasts (p-PNA) and mitochondria (m-PNA), diluted into a master mix at 2.5 μM according to manufacturer’s instructions. PCR Reactions (per sample) consisted of 12.5 μL HotStart Taq 2x Polymerase Master Mix (New England Biolabs), 2 μL forward primer master mix, 2 μL reverse primer master mix, 2.5 μL of PNA master mix, 1 μL of sterile water, and 5 μL of eluted DNA template. The first amplification PCR protocol was 95°C for 45 seconds; 20 cycles of 95°C for 15 seconds, 78°C for 10 seconds, 60°C for 45 seconds, and 72°C for 45 seconds; and finally hold at 4°C. Amplified DNA was purified using AMPure XP beads (Beckman Coulter) according to the manufacturer’s protocol.

The second-step PCR was used to add Illumina barcodes (for multiplexing) and sequencing adaptors, using Illumina Nextera Index adaptors N701-N729 and S502-S522 (24 x 16 = 384 total combinations). Index combinations for each well were pre-mixed in a separate 96-well plate to 2.5 μM per primer. PCR reactions consisted of (per sample) 12.5 μL HotStart Taq 2x Polymerase Master Mix (New England Biolabs), 2.5 μL of PNA master mix, 5 μL dual-index primer mix (2.5 μM each), and 5 μL of eluted DNA template. The indexing PCR protocol was 95°C for 45 seconds; 25 cycles of 95°C for 15 seconds, 78°C for 10 seconds, 60°C for 45 seconds, and 72°C for 45seconds; 68° for 5 minutes; and finally hold at 4°C. The resulting PCR products were purified with AMPure XP beads (Beckman Coulter) according to the manufacturer’s protocol.

The final sequencing library was sent to the UGA Genomics and Bioinformatics Core for quality control and sequencing with paired-end 300 sequencing on an Illumina MiSeq.

### Read Processing

The primer sequences from the raw data were excised using Cutadapt (Martin 2011), which also filtered out reads with a Phred quality score below 20. Read quality was visually assessed with MultiQC (Ewels et al., 2016). QIIME2 version 2022.11 (Bolyen et al., 2019) was used to process trimmed sequences. The DADA2 (Callahan et al., 2016) plugin was used to analyze read quality profiles, and sequences with an average quality score less than 30 were removed. Paired-read joining, generation of representative sequences, alignment of the representative sequences, and chimera checking were all carried out using the DADA2 plugin in QIIME2. The taxonomic classification of amplicon sequence variants (ASVs) was assigned using the SILVA 138-99-nb pretrained classifier (Quast et al., 2012). Taxa without phylum identification were discarded, as were chloroplasts and mitochondria. Samples with less than 500 reads were also removed. A total of 717 putative contaminant ASVs (as determined by their presence in blank samples) were eliminated using the R package decontam v1.10.0, as described by (Davis et al., 2018).

### Diversity metrics

We assessed alpha diversity using three widely used metrics: observed ASVs, Shannon entropy, and Faith’s phylogenetic distance (PD). Datasets were rarefied to the approximate minimum reads per sample using the phyloseq package (McMurdie & Holmes, 2013), resulting in 2000, 3000, 1000, and 500 for soil, rhizosphere, root, and stalk, respectively. Alpha diversity scores were tested for normality using the Shapiro-Wilkes test; since the data were not normally distributed, significance testing was conducted with a Kruskal-Wallis test followed by Dunn’s post-hoc test (with Bonferroni correction) through the FSA package (Ogle et al., 2022). A general linear model was used to check for associations between alpha diversity and environmental variables.

The beta-diversity of bacterial communities in each tissue type was assessed by performing Principal Coordinates Analysis (PCoA) with Weighted UniFrac distance matrices (Lozupone et al., 2011). The bacterial microbiome dissimilarity distance metrics were also analyzed by the multivariate permutation analysis of variance (PERMANOVA), together with the nonparametric statistical Adonis test (vegan package) to establish the homogeneity of dispersion. The PERMANOVA test was performed using 999 permutations. To test whether microbial community similarity (or dissimilarity) is related to geographic distance between samples, we used Spearman rank correlation of distances between samples by weighted unifrac and by geographic distance.

Sampling noise was estimated by comparing alpha and beta diversities for pairs of plant in R using the MicrobiomeStat package (Chen et al., 2022).

### Environmental descriptors and their association with the microbiome communities

Environmental variables for this study fall into three classes: (1) **Geography**: Latitude (LAT), Longitude (LONG). (2) **Soil Properties**: Lime Buffer Capacity (LBC), Soil Ph (pH), % Base Saturation (BSP), Cation Exchange Capacity (CEC), Calcium (ppm) (Ca), Cadmium (ppm) (Cd), Chromium (ppm) (Cr), Copper (ppm) (Cu), Iron (ppm) (Fe), Potassium (ppm) (K), Magnesium (ppm) (Mg), Manganese (ppm) (Mn), Molybdenum (ppm) (Mo), Sodium (ppm) (Na), Nickel (ppm) (Ni), Phosphorus (ppm) (P), Lead (ppm) (Pb), Zinc (ppm) (Zn), % Organic Matter (OM), % Nitrogen (Np). (3) **Weather**: Precipitation Probability (PP), Precipitation Intensity (PI), High Temperature (HT), Low Temperature (LT), Dew Point (DP), Humidity (HUM), Wind Speed (WS), UV Index (UVI). To assess the correlation of beta diversity with environmental variables, each ASV table was transformed to relative abundance and weighted UniFrac distances were calculated using the rbiom (Smith, 2023) and vegan (Dixon, 2003) R packages. To determine whether the difference in community composition among locations was spatially correlated, we used the R package “geosphere” to calculate the Haversine distances (distance between two points on a sphere) given latitude and longitude (Hijmans et al., 2015). Mantel test was used to examine the strength and significance of the association. Afterwards, the relationship between environmental factors and microbial beta diversity was identified. We used canonical correspondence analysis (CCA) combined with ANOVA based on the decorana() function, which is a diagnostic tool to choose between CCA and RDA based on the response patterns of the species data. For CCA, Variance inflation factors (VIF) were computed to assess multicollinearity. Variables with VIF > 10 were considered collinear and removed to reduce redundancy. CCA models were fitted separately for each variable group using the cca() function in the vegan package. Forward selection, which identifies the most influential predictors within each variable group, was conducted using the ordistep() function in the vegan package. Model significance and the marginal (by-term) effects of individual variables were evaluated with permutation ANOVA (999 permutations). Only variables with p < 0.05 were displayed on the CCA biplot.

To investigate the correlation between various Taxa with the environmental factors, ASVs were correlated with environmental variables using Spearman correlation in the microeco package in R (Liu et al., 2021). ASVs were converted to relative abundance and then agglomerated to the class level because aggregation at the class level improved interpretability and revealed consistent ecological patterns, while grouping was done using growing degree days (GDD) as a factor. P-values were corrected for multiple testing using the Benjamini & Hochberg (False Discovery Rate) approach (Benjamini & Hochberg, 1995).

### Differential abundance analysis of taxa

To identify the bacterial ASVs enriched within the different tissue types and GDDs, we used a negative binomial model to test for differential abundance using the package DESeq2 (Love et al., 2014). To minimize the risk of false positives, we filtered the adjusted p-values so that only ASVs with p-values less than 0.001 are highlighted in this study.

### Network co-occurrence analysis

ASV co-occurrence networks were constructed using only those ASVs with relative proportions greater than 0.01%. Spearman’s Rho between the pairwise ASVs matrices were constructed using the “microeco” package in R (Liu et al., 2021). The false discovery rate (FDR) was used to correct p-values to account for multiple testing (Benjamini & Hochberg, 1995). Co-occurrence was considered robust if the absolute value of the Spearman correlation coefficient was either equal or greater than 0.6 and statistically significant if p-values < 0.05. The co-occurrence networks were assessed using the R package igraph 1.2.6 (Csardi, 2013) and visualized using Gephi 0.9.2 (Bastian et al., 2009). In the network, nodes represented ASVs, whereas edges represented the correlation between nodes; network layout was based on an undirected network with the OpenOrd layout. To evaluate the significant differences between the constructed network and its corresponding random network, the Erdos–Renyi model (Erdős & Rényi, 1959) was used to generate random networks with the same size (the same number of nodes and edges) as the real network using the “microeco” R package.

### Metagenomic Prediction of microbial communities

The metagenomic prediction of each sample type was performed with Phylogenetic Investigation of Communities by Reconstruction of Unobserved States (PICRUSt 2.6.3) (Douglas et al., 2020). This tool facilitated the prediction of the metagenome by utilizing the data derived from the 16S ribosomal RNA amplicon datasets for each sample type. Subsequently, the functional potential was performed with the q2-picrust2 implementation tool, as developed by G.M. Douglas (available at https://github.com/gavinmdouglas/q2-picrust2/). The metagenomic predictions obtained were then subjected to functional annotation, employing the Kyoto Encyclopedia of Genes and Genomes (KEGG) pathways (Kanehisa et al., 2017) framework using ggpicrust2 in R (Yang et al., 2023). This package was also used for abundance conversion, pathway annotation, differential abundance (DA) analysis, and DA results visualization. Within each compartment, functions that changed abundance between the two timepoints were calculated by DESeq2 differential abundance analysis (p < 0.05, FDR-adjusted), with KEGG ortholog (KO) abundances first converted to KEGG pathway abundances using the ko2kegg_abundance function.

## Results

### Sample collection and sequencing

A total of 35 fields at 16 locations around the state of Georgia were sampled at two timepoints during the summer of 2018 (**Figure 1A**); timepoints were chosen to coincide with the initiation of the ear (∼600 GDDs) and flowering (∼1400 GDDs), respectively. At each field, two plants were selected and the stalk, root, rhizosphere, and bulk soil samples collected on dry ice (**Figure 1B**). DNA was extracted from each, and bacterial 16s sequences amplified following using Earth Microbiome Project primers (Caporaso et al., 2012). After removing samples with failed amplification or insufficient DNA, a total of 439 samples plus 12 blanks (3 per sample type) were sequenced.

Amplicon sequence variants (ASVs) were called using DADA2 in QIIME2 (Bolyen et al., 2019; Callahan et al., 2016). Probable contaminant ASVs removed with the R package Decontam, which improves the quality of metagenomic data by identifying and removing contaminant ASVs was used; a total of 717 ASVs (2.1%) contaminants found in all the sample types were removed (**Supplementary Figure 1**). Rhizosphere samples had the highest average sequencing depth while stalk samples had the lowest (**Supplementary Figure 1, Table 1**). Collectively, our final dataset comprised 334 samples and 38,418 Amplicon Sequence Variants (ASVs).

**Table 1:** The sequencing depth result and other related information for the respective samples.

| **Samples** | **Number of samples** | **Number of reads** | **Median read depth** | **Number of ASVs** |
|---|---|---|---|---|
| **Root** | 90 | 788,170 | 4881 | 7282 |
| **Stalk** | 66 | 310,016 | 2602.5 | 1608 |
| **Soil** | 68 | 1,009,206 | 10,664.5 | 11,615 |
| **Rhizosphere** | 110 | 1,547,065 | 12,978 | 17,913 |

### Bacterial communities across maize compartments

Proteobacteria became more abundant when moving from soil (18.7%) to stalk (52.3%) (**Figure 2**). Proteobacteria, alongside Chloroflexi and Acidobacteriota were most abundant in the soil (19.4%, and 19.0% respectively). On the other hand, Actinobacteria was most abundant in the root (20.79%) and least abundant in the stalk (7.97%). The abundance of Proteobacteria in the rhizosphere, root, and soil declined by 0.09%, 0.12%, and 0.09% respectively at least 1.6% between 600 GDDs and 1400 GDDs, but the inverse was observed in the stalk (**Figure 2**). In the rhizosphere, the abundance of Acidobacteriota and Chloroflexi increased from 600 GDDs to 1400 GDDs, while Bacteriodota and Actinobacteriota declined in abundance. A similar trend was observed in root interior samples as the abundance of Acidobacteriota and Chloroflexi increased by 0.01% and 0.78% respectively with Actinobacteriota and Bacteriodota decreasing in abundance across sample points by 0.15% and 0.15% respectively. In the soil, the abundance of Acidobacteriota and Armatimonadota declined, whereas Actinobacteriota and Chloroflexi slightly increased between samplings. Lastly, in the stalk, the abundance of Acidobacteriota, Actinobacteriota, and Chloroflexi declined 0.57%, 0.5%, and 0.78%, whereas Bacteroidota increased from 11.4% to 28.69% across samplings.

**Figure 2:**
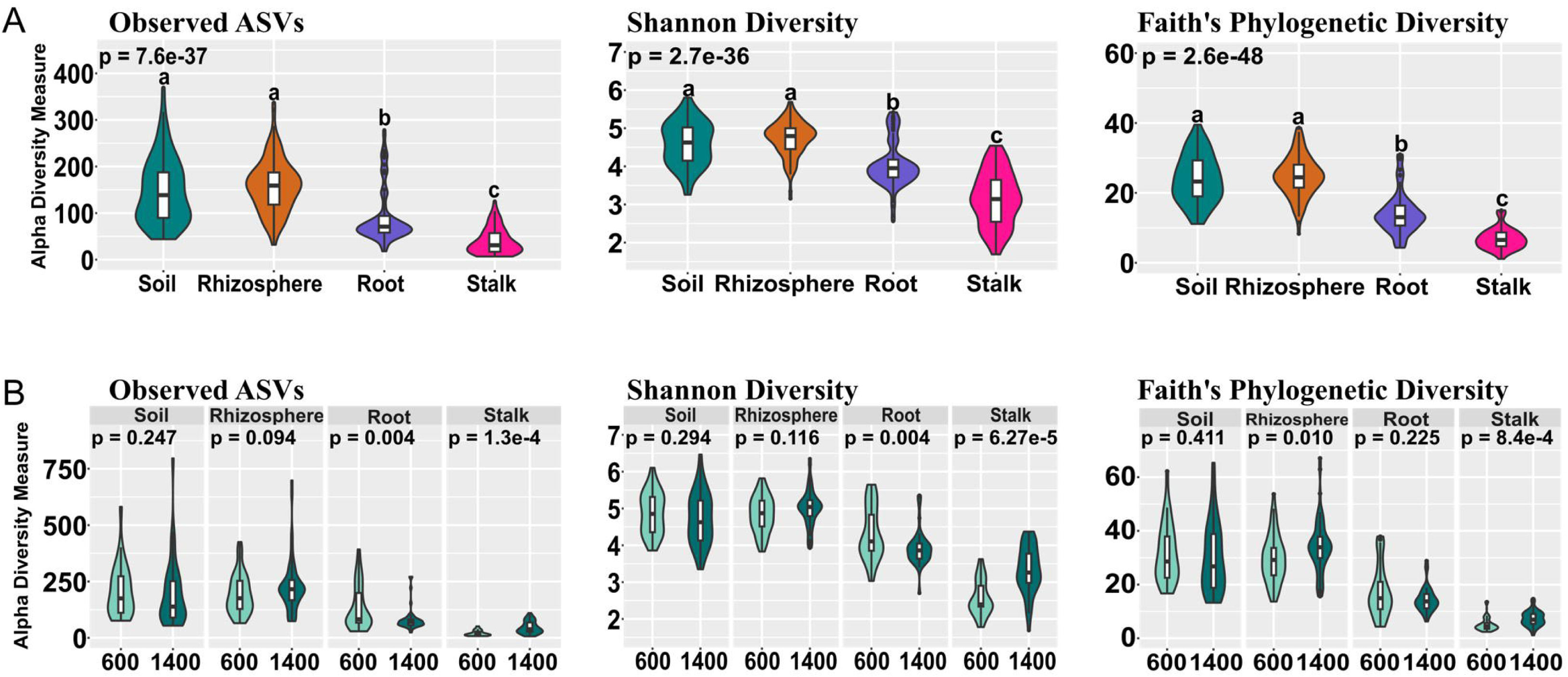

### Alpha Diversity

Rhizosphere and soil samples had higher alpha (within-sample) diversity than roots and stalks across the various metrics (**Figure 3A**). Post-hoc tests show that soil and the rhizosphere did not show significant changes in alpha diversity between sample times for Observed and Shannon diversity, though the rhizosphere showed was significant for Faith’s PD (**Figure 3B**). Root samples, however, showed a decrease in diversity for Observed and Shannon diversity metrices but not for Faith PD while stalk samples showed a highly significant increase across the three metrics (**Figure 3B**). Alpha diversity also varied widely by location (**Supplementary Figure 2**). In the rhizosphere, significant variation was observed for Observed ASVs (p = 0.042), Shannon diversity (p = 0.028), and Faith’s PD (p = 0.049), indicating spatial heterogeneity in microbial community richness and phylogenetic structure. Root samples exhibited a significant difference only in Faith’s Phylogenetic Diversity (p = 0.007), suggesting that community phylogenetic breadth varied more than general taxonomic richness or evenness. Stalk samples showed significant variation in Observed ASVs (p = 0.043) but not in Shannon or Faith’s diversity indices, reflecting richness differences without pronounced changes in evenness or phylogenetic diversity. In contrast, soil samples showed no significant differences across locations for any of the alpha diversity metrics, indicating relatively uniform diversity in this compartment despite significant differences in soil composition.

**Figure 3:**
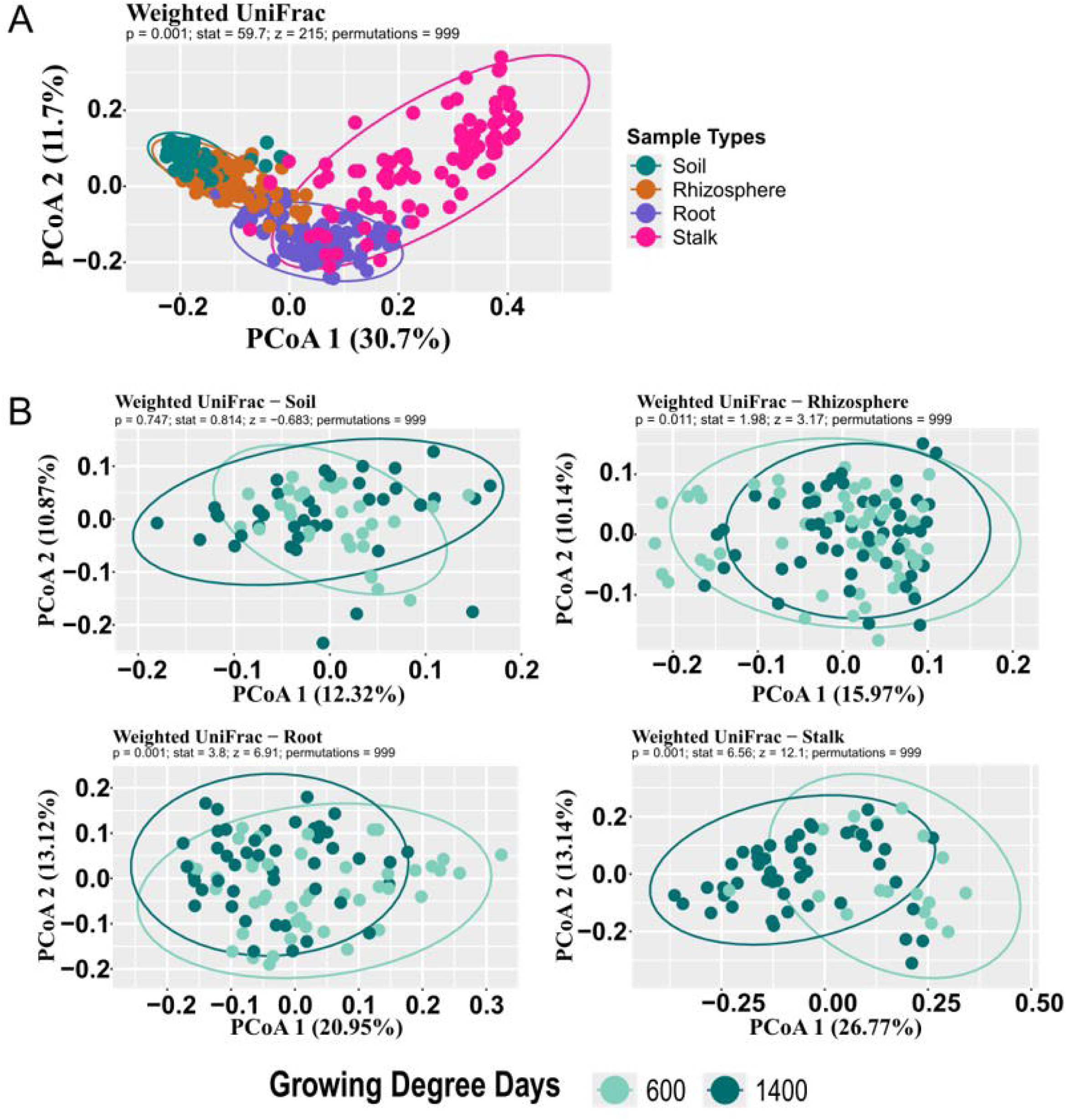

### Beta Diversity

Beta (between-sample) diversity was calculated with weighted UniFrac and revealed sample separation based on sample type, with each forming a distinct cluster (**Figure 4A**). Sample types were significantly different from each (PERMANOVA p=0.001, 999 permutations). Within each sample type, samples from different times were also significantly different, except in soil samples (**Figure 4B**). At 600 GDDs, the geographic distance was significantly but weakly correlated with UniFrac distance at individual sample level for soil (Mantel’s r = 0.150, p = 0.023) rhizosphere (Mantel’s r = 0.267, p = 0.001) and root (Mantel’s r = 0.095, p = 0.023) samples but not for stalk (Mantel’s r = 0.063, p = 0.305) (**Supplementary Figure 3**). At 1400 GDDs, soil (Mantel’s r = 0.267, p = 0.002) and rhizosphere (Mantel’s r = 0.14, p = 0.002), and root (Mantel’s r = 0.093, p = 0.028)showed significant correlation whereas stalk (Mantel’s r = 0.003, p = 0.467) showed no significant spatial relationship.

**Figure 4:**
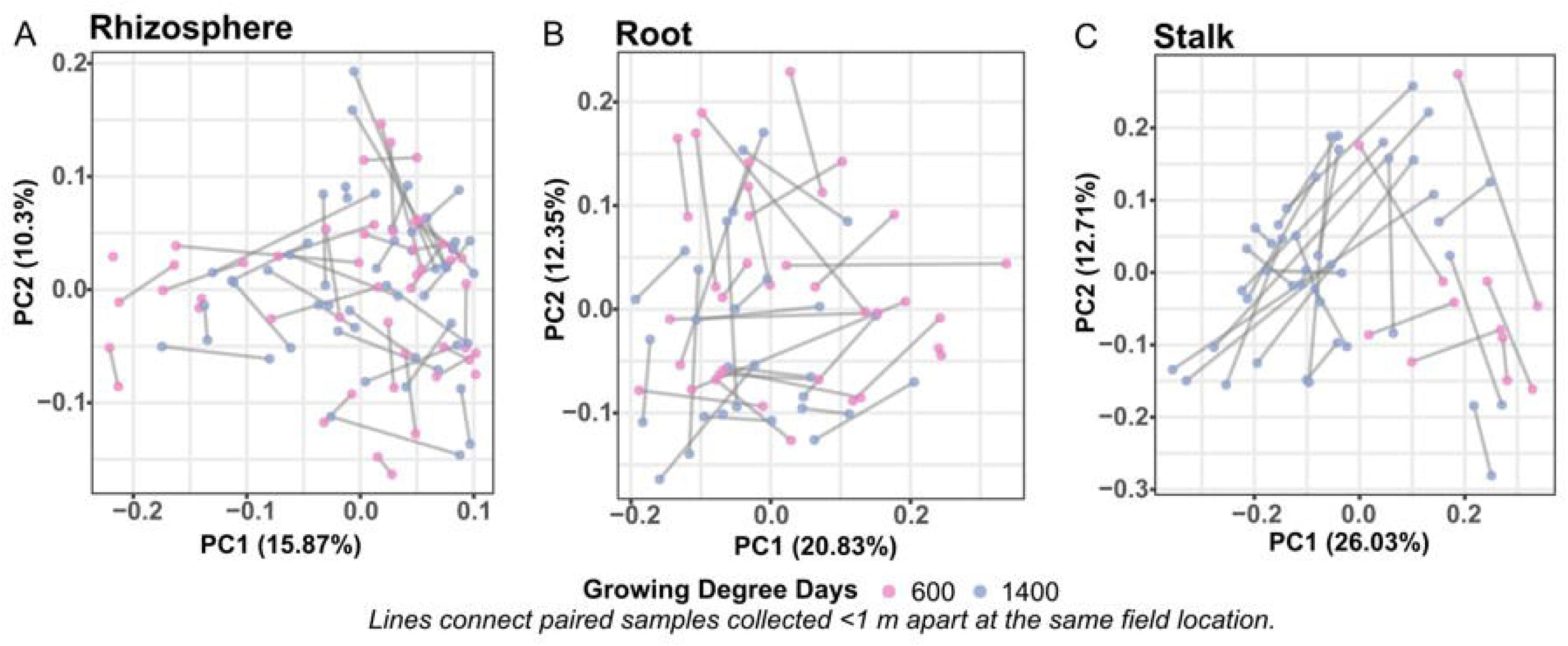

### Estimating the sampling error of the microbiome

At each location, two plants ∼1 meter apart were taken to sample the plant-associated microbiome (rhizosphere, root, stalk). Since all sampled plants were hybrids, each pair is genetically identical and so represents the (presumably) minimum variation possible within a field. As such, they can be used to determine the intrinsic noise in sampling these microbiomes. After quality filtering, we were left with 32 root pairs, 49 rhizosphere pairs, and 28 stalk pairs. Alpha diversity metrics exhibited compartment-specific patterns of within-pair variability (**Table 2**). Rhizosphere communities showed the lowest variability across most metrics, with Shannon diversity exhibiting a mean coefficient of variation (CV) of 6.7% ± 5.1% and Simpson diversity showing exceptional consistency (CV = 0.36% ± 0.34%). Root and stalk compartments showed slightly higher but still moderate Shannon diversity variability (CV = 10.1% ± 9.5% and 11.3% ± 10.4%, respectively). Pielou’s evenness index remained stable across all compartments (CV < 3.3%). In contrast, metrics based on taxonomic richness revealed prominent within-pair variation across all compartments. For observed species richness, root samples recorded a mean CV of 34.5% ± 29.3% (mean relative difference (MRD) of 59.2% ± 78.7%). Stalk also recorded a CV of 34% ± 24.1%, and rhizosphere shows a CV of 30.2% ± 21.8%. Chao1 richness for root samples shows a mean CV of 36.3% ± 31.5% (MRD: 65.1% ± 95.7%). The high variability in richness estimates likely reflects the combined effects of genuine microhabitat heterogeneity at sub-meter scales, stochastic sampling of rare taxa, and inherent uncertainty in richness estimation from rarefied data. In contrast, evenness-based metrics showed substantially lower within-pair variability across all compartments (Shannon CV: 6.7–11.3%; Simpson CV: ≤1%; Pielou’s evenness CV: ≤3.3%), indicating that the dominant community structure is highly reproducible at sub-meter scales. Taken together, these results suggest that while the total number of detectable taxa varies between nearby samples due to stochastic rare-taxon sampling, the overall community composition and relative abundance of dominant members are stable within sampling sites. Across all compartments, weighted UniFrac distances among sample pairs are significantly more similar to each than among random samples (**Figure 5; Supplementary Figure 4**). Rhizosphere pairs were the most similar to each other while stalks were the least (**Table 2**), though all were significant at p < 0.01.

**Figure 5:**
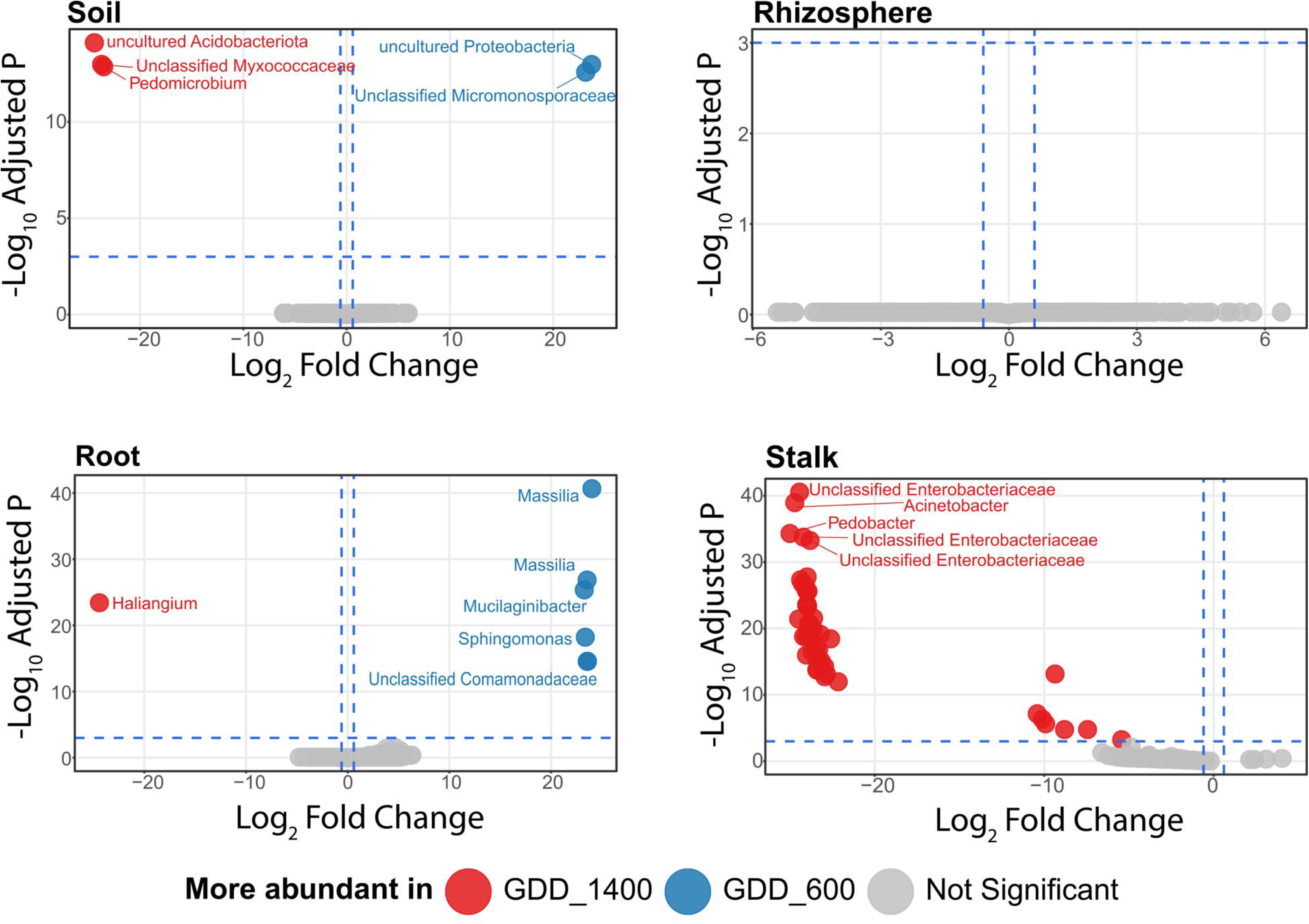

**Table 2:**
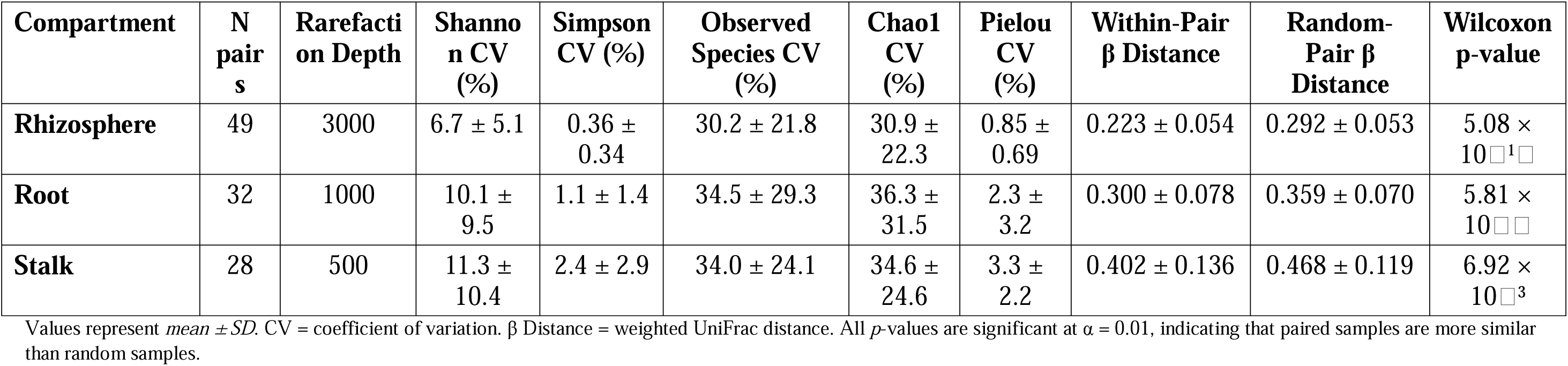
Summary of Diversity Metrics and β Distances Across Compartments.

### Differentially abundant microbes across compartments

Differential abundance analysis was conducted at the ASV level using DESeq2 based on variables of interest (sample types and timepoints). Following statistical testing, taxonomic annotations were merged with the ASV-level results (**Supplemental Table 1**), and Genus names were used solely for labeling and interpretation of the most significantly differentially abundant ASVs in the volcano plots. Volcano plots of differential abundance (**Figure 6**) reveal that soil communities show a relatively balanced pattern, with significant significant abundant ASVs enriched in both timepoints, potentially indicating community succession. The reverse was observed for root, where Haliangium is notably the only highly significant abundant genus more abundant at the later timepoint (1400 GDDs), whereas many significant ASVs particularly Massilia, Mucilaginibacter, and Sphingomonas became more abundant at 600 GDDs. Stalk communities, meanwhile, had Unclassified Enterobacteriaceae, Pedobacter, Acinetobacter genera significantly abundant at the second timepoint. Interestingly for rhizosphere communities, there were no significantly differentially abundant genera at both timepoints. These contrasting patterns of significantly differential abundance across the different compartments highlights different trajectories for each community across developmental stages.

**Figure 6:**
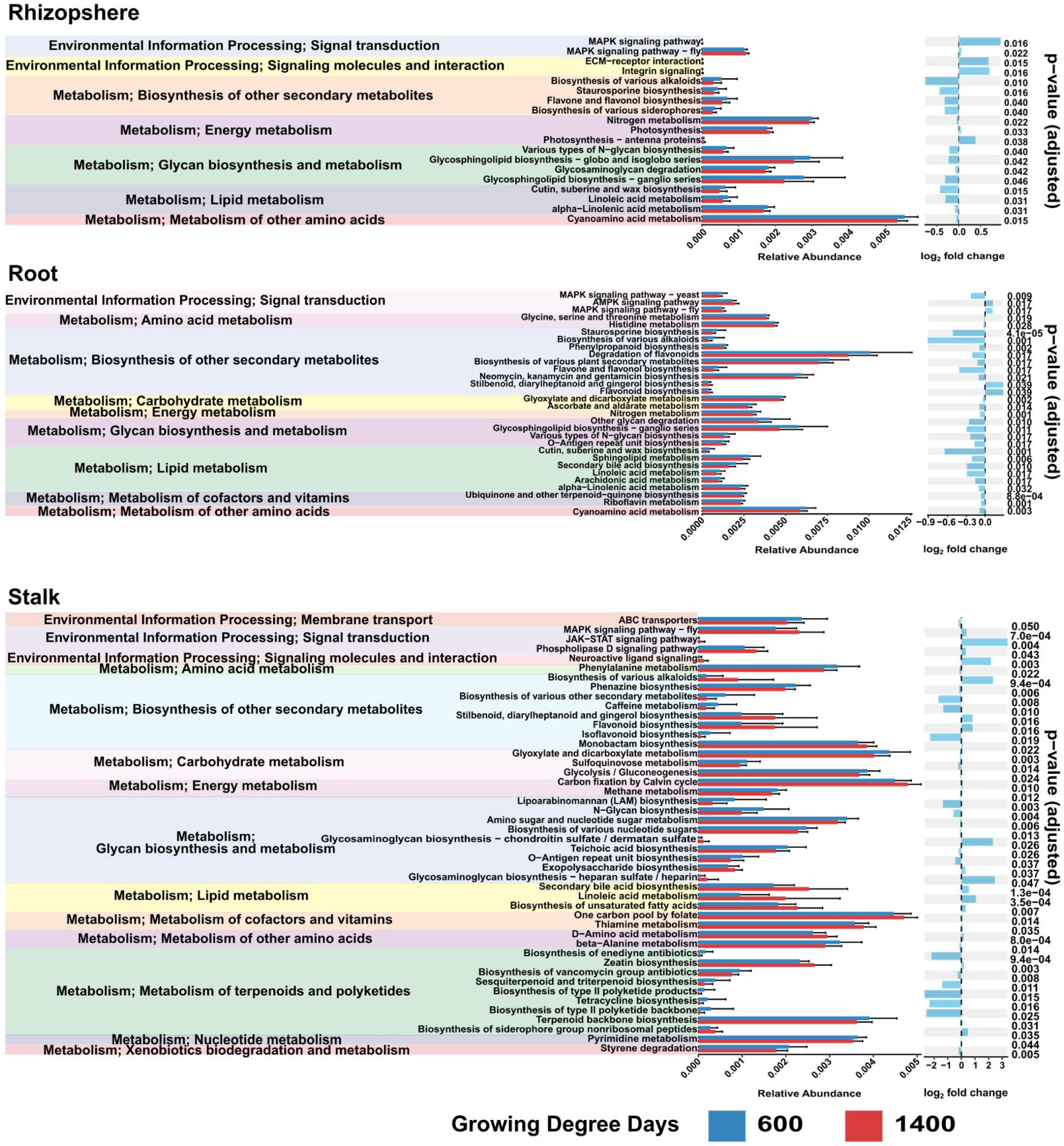

### Functional capacity of microbial communities

Functional capacity of each community was estimated with PICRUSt2 (Douglas et al., 2020). First, we uncovered the gene functions predicted for each sample type across the two GDD. Rhizosphere, stalk, and root bacterial community showed statistically significant pathways, whereas there were none for soil communities (**Figure 7**). In the rhizosphere, communities with functions related to metabolism such as nitrogen, alpha-linoleic acid, cyanoamino acid metabolism, biosynthesis of various alkaloids, flavone and flavanol biosynthesis, and glycosaminoglycan degradation were more enriched at 600 GDDs than at 1400 GDDs. However, at the later stage, communities showed greater relative abundance of the MAPK signaling pathway. In the root, clear functional divergence was also observed between the two growth stages. Cyanoamino acid metabolism, degradation of flavonoids, biosynthesis of various plant metabolites, and glycosphingolipid biosynthesis was relatively higher at 600 GDDs, whereas MAPK and AMPK signaling pathways increased at 1400. In the stalk, functional enrichment at 600 GDDs was dominated by genetic information processing and energy metabolism; specifically, sulfur relay systems, ABC transporter, quorum sensing, biosynthesis of various other secondary metabolites, glyoxylate and dicarboxylate metabolism were more abundant. The stalk communities at 1400 GDDs were more enriched in MAPK signaling pathway; stilbenoid, diarylheptanoid and gingerol biosynthesis; and biosynthesis of zeatin, monobactam, various alkaloids, flavonoids, etc.

**Figure 7:**
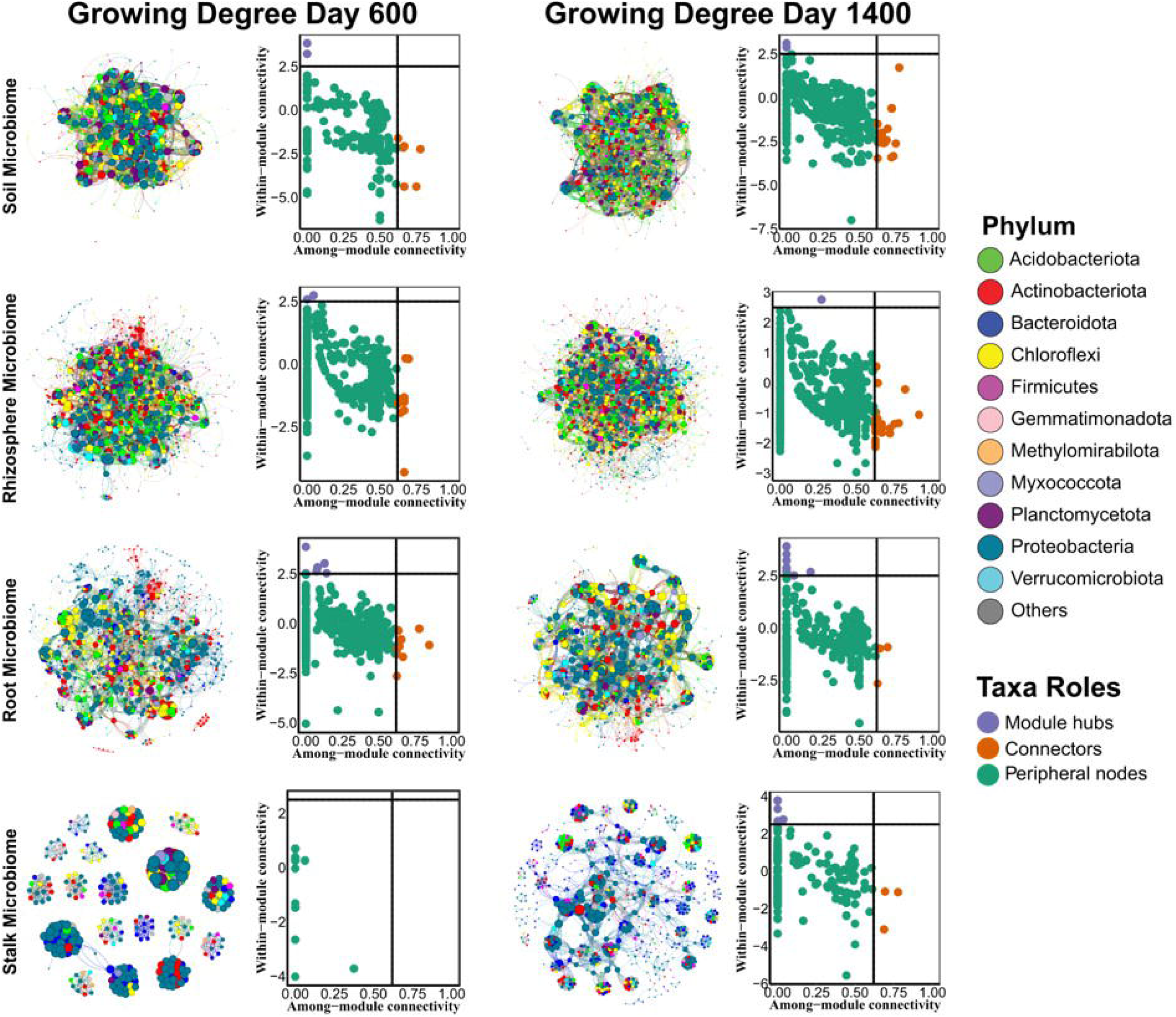

### Network co-occurrence analysis for bacterial communities

To identify potential interactions among bacteria, we constructed a co-occurrence network for each sample type. In terms of the network-level topological features, both the average path length and average clustering coefficient were greater than in random networks across the GDDs (Table 5; Supplemental Table 2). At 600 GDDs, the stalk microbiome shows the highest clustering coefficient (0.994), indicating a more tightly interconnected microbial network than those observed in soil (0.735), root (0.699), and rhizosphere (0.654); this ordering is preserved at 1400 GDDs, except that root and soil (the middle two) switched places (**Table 3**). Average path length was lowest in stalk at 600 GDDs (1.322), indicating a compact network and efficiently connected microbial interactions; however, by 1400 GDDs, stalks had the highest path length (4.149), whereas the rank order of the other three samples remained the same (soil < rhizosphere < root) (**Table 3**).

**Table 3:** Topological properties of stalk, rhizosphere, soil, and root bacterial communities across the two growing degree days.

| Real Networks |  |  |  |  |  |  |  | Random Network<br>(Erdos-Reyni) |  |
|---|---|---|---|---|---|---|---|---|---|
| Growing Degree Days | Tissue Types | Nodes | Edges | Average degree | Network Diameter | Average clustering coefficient | Average path length | Average clustering coefficient | Average path length |
| **600** | **Soil** | 1374 | 36,027 | 52.441 | 6 | 0.735<sup>**</sup> | 2.346<sup>**</sup> | 0.0381<br>± 0.000226 | 2.091<br>± 0.000404 |
|  | **Rhizosphere** | 1439 | 16,711 | 23.226 | 9 | 0.654<sup>**</sup> | 3.109<sup>**</sup> | 0.016175<br>± 0.000351 | 2.661264<br>±0.000341 |
|  | **Root** | 984 | 8558 | 17.394 | 8 | 0.699<sup>**</sup> | 3.115<sup>**</sup> | 0.017624<br>± 0.000684 | 2.717017<br>± 0.000673 |
|  | **Stalk** | 317 | 3398 | 21.438 | 4 | 0.994<sup>**</sup> | 1.322<sup>**</sup> | 0.067943<br>± 0.001539 | 2.150175<br>± 0.001829 |
| **1400** | **Soil** | 1596 | 31,653 | 39.665 | 6 | 0.743<sup>**</sup> | 2.567<sup>**</sup> | 0.024871<br>± 0.000221 | 2.338867<br>± 0.000325 |
|  | **Rhizosphere** | 1404 | 13,630 | 19.416 | 8 | 0.608<sup>**</sup> | 3.097<sup>**</sup> | 0.013889<br>± 0.00038 | 2.75279<br>± 0.000447 |
|  | **Root** | 1177 | 15,303 | 26.003 | 8 | 0.760<sup>**</sup> | 3.147<sup>**</sup> | 0.022065<br>± 0.000385 | 2.528357<br>± 0.000407 |
|  | **Stalk** | 775 | 6481 | 16.725 | 11 | 0.854<sup>**</sup> | 4.149<sup>**</sup> | 0.021559<br>± 0.000694 | 2.668636<br>± 0.000764 |

Observed network metrics were compared with 999 Erdős–Rényi random networks generated using the random\_network() function in the *microeco* package. Asterisks indicate significant differences from the random networks (*p* < 0.05, **p** < 0.01, ***p*** < 0.001).

In addition to overall network shape, the specific network attributes of individual ASVs can provide insight into their ecological function (**Figure 8**). ASVs were classified based on the within-module connectivity (Zi) and among-module connectivity (Pi) into three categories. Peripheral ASVs have low Zi and low Pi, indicating few links to other ASVs; the majority of ASVs fell into this category across all sample types. Module hubs have a high Zi and a low Pi, indicating a substantial number of connections within their own modules but few outsides of it. Connectors typified by a low Zi and a high Pi, establish linkages across multiple modules. For our analysis, nodes with Pi ≥ 0.62 were classified as connectors, one with Zi ≥ 2.5 were classified as modules, and the rest were peripheral. (No ASVs had both Pi ≥ 0.62 and Zi ≥ 2.5 in any network.) With these cutoffs, the soil, rhizosphere, root, and stalk networks have 5, 4, 15, and 0 module hubs respectively (**Figure 8; Supplemental Table 2**) at 600 GDDS, and 3, 6, 16, and 9 module hubs respectively at 1400 GDDs. For connectors, the soil, rhizosphere, root and stalk networks had 6, 53, 28, and 0 connectors 600 GDDs and 36, 54, 3, 3, respectively, at 1400 GDDs.

**Figure 8:**
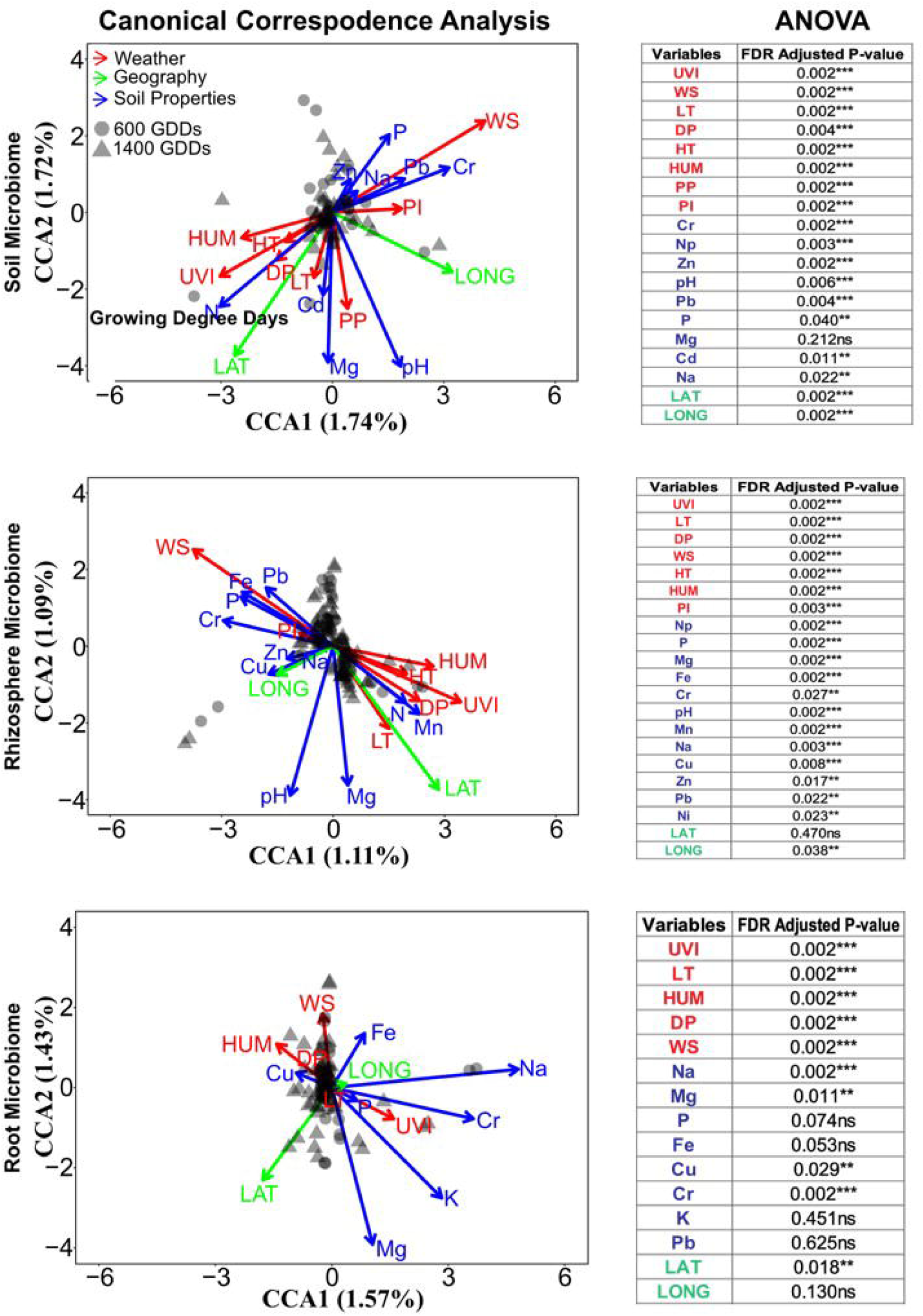

Across both timepoints, several taxa from distinct phyla served as key network components (**Figure 8; Supplemental Table 2**). Proteobacteria were the predominant module hubs across all networks, with Acidobacteriota additionally serving this role in soil and rhizosphere networks, and Actinobacteriota in the root network. Similarly, Proteobacteria were the predominant connectors across all networks, with Acidobacteria additionally prominent in soil networks, Actinobacteriota in the rhizosphere, and Chloroflexi in both soil and root networks. Proteobacteria thus consistently emerged as a central phylum across all maize compartments and developmental stages, with members serving as both major module hubs and connectors.

### Environmental impact on the microbiome communities

Weather data for each location (humidity, precipitation, temperature, etc.) was collected using the darksky R package (Rudis 2017) and combined with physical characteristics from soil samples. Linear regression between these factors and alpha diversity (Shannon diversity; Faith’s PD) showed that most metrics were not associated with diversity, and those that were varied depending on the compartment (Supplemental **Table 3**). Based on the p-value for bulk soil, both metrics were associated with humidity, while rhizosphere diversity correlated with % nitrogen for Shannon diversity. In the root endosphere, only Faith’s PD correlated with any metrics (sodium, lime buffer capacity). The stalk community showed the most environmental associations, with one or both metrics significantly associated with low temperature, dewpoint, latitude, cadmium, copper, sodium, and phosphorus. However, with multiple testing correction based on false discovery rate (FDR), only latitude was significantly associated with Shannon for stalk samples.

We further assessed the relationship between community makeup and environmental factors with canonical correspondence analysis (CCA), a multivariate statistical technique used to relate species’ abundance and environmental variables (ter Braak and Verdonschot, 1995), with multiple testing correction based on false discovery rate (FDR). CCA identified significant associations for soil, root, and rhizosphere bacterial communities, but not for stalk bacterial communities. Both low temperature and wind speed showed consistent significant relationships across the three associated bacterial communities (**Figure 9**). Soil communities were also associated with UV Index, Humidity, Wind Speed, Dew Point, Precipitation Intensity, Precipitation Probability, High Temperature, Low Temperature, Latitude, and Longitude, as well as several soil chemical properties (Chromium, Zinc, %Nitrogen, Phosphorus, Cadmium, Sodium, pH, and Lead). Rhizosphere communities were associated with UV Index, Low Temperature, Dew Point, Wind Speed, High Temperature, Humidity, Precipitation Intensity, %Nitrogen, Phosphorus, Magnesium, Iron, Chromium, pH, Manganese, Sodium, Copper, Zinc, Lead, and Longitude, while root communities were associated with UV Index, Humidity, Low Temperature, Dew Point, Wind Speed, Sodium, Magnesium, Iron, Copper, Chromium, and Latitude.

**Figure 9:**
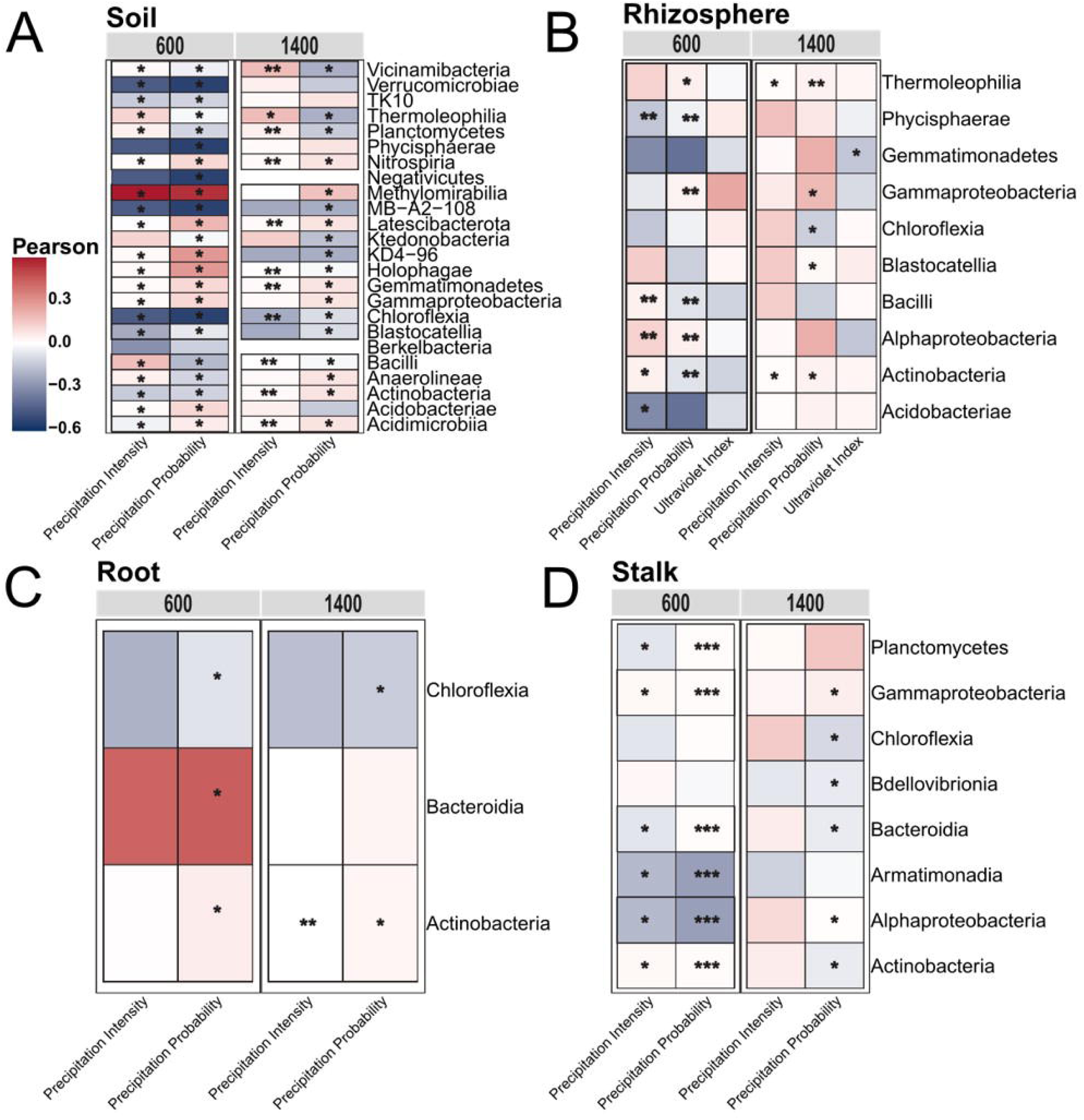

At a finer scale, we performed Pearson correlation analysis between environmental variables and the relative abundances of individual taxonomic class to identify potential drivers of specific bacterial taxa. Precipitation (both probability and intensity) were the main factors that correlated with class abundance across the four sample types (**Figure 10**). However, the significance of this correlation varied across the two sample times in the same bacterial community. For instance, for the root community, precipitation probability showed a significant positive correlation with Bacteroidia at 600 GDDs but was nonsignificant at 1400 GDDs. In other cases, the correlation between an environmental variable and a phylum varied across different bacterial communities at the same timepoint. For instance, at 1400 GDDs, precipitation probability showed a significant positive correlation with Actinobacteria in the root community but a significant negative correlation with it in stalk communities. Similar patterns held for individual bacterial classes. For example, while Na, K, and cation exchange capacity (CEC) showed a significant positive correlation with the class Thermoleophilia at 600 GDDs, they were negatively correlated at 1400 GDDs (**Supplementary figure 5)**. This pattern was also observed for rhizosphere, root and stalk communities. Overall, the significant correlation of environmental variables and soil chemical properties variables on class was highly variable, based on the microbiome compartment and sampling time.

## Discussion

The maize microbiome has been extensively investigated (Favela et al., 2021; Jiang et al., 2024; Kong et al., 2020; Rüger et al., 2021; Sanchez-Garnica et al., 2024; Xiong et al., 2021a; Zhang et al., 2024c). However, these analyses are usually performed under conditions that do not reflect the actual growing conditions of farmers. In this study, we analyzed the maize bacterial communities in four compartments (soil, rhizosphere, root interior, shoot interior) from commercial hybrids grown in actual production systems and at two critical time periods (600 and 1400 growing-degree days, approximately the time of ear formation and flowering, respectively). These results shed light on the variation and evolution of the maize microbiome under actual production conditions and with commercially relevant varieties.

### Microbial community composition and abundance

Across sample types, the phyla Proteobacteria, Chloroflexi, Actinobacteriota, Planctomycetota, and Acidobacteriota were most observed, similar to other plant and soil associated microbiomes (Ahkami et al., 2017; Nutaratat et al., 2024; Pu et al., 2022; Zeng et al., 2016). Proteobacteria were the most abundant phyla in roots and stalks, with high abundance also in rhizosphere and soil samples. Members of this phylum contributes to various plant beneficial functions such as nitrification, nitrogen fixation, and methylotrophy (Atamna Ismaeel et al., 2012; Watanabe et al., 2016). Acidobacteriota was dominant in the soil, matching prior studies (Lauber et al., 2009; Nemergut et al., 2010; Roesch et al., 2007), and lower in roots and stalks, possibly because bacteria in this phylum tend to be oligotrophic and favor low-nutrient environments. Chloroflexi was also prevalent in soil and rhizosphere; this phylum has been implicated in soil physical disturbance, which in turn impacts microscale topography, nitrogen availability, and a shift in the retention of moisture (Kuske et al., 2012). We consistently detected Actinobacteriota across all sample types, which support findings that they are widespread in both plant endospheres and soil environments (Ahkami et al., 2017; Yadav et al., 2021), including maize stalks (Cobo-Díaz et al., 2019). It is also noteworthy that timepoint (growing degree days) influenced the relative abundance of these phyla in a sample type-specific manner. For instance, the phyla Chloroflexi abundance increased as we moved from 600 GDDs to 1400 GDDs, but the reverse was the case for the same phyla in the root.

### Drivers of diversity across microbial communities

Several factors appear to be driving overall microbial diversity. The largest of these is sample type, with the soil, root and rhizosphere (belowground) sampling having higher alpha diversity than stalks (above ground). However, between soil and the rhizosphere, there was no significant difference. The same pattern has been seen previously in maize (Schultz et al., 2023)and other plants including Arabidopsis (*Arabidopsis thaliana*), soybean (*Glycine max*), rice (*Oryza sativa*), tomato (*Solanum lycopersicum*), Agave (*Agave spp*.) and some of its relatives (Delmotte et al., 2009; Dong et al., 2019; Knief et al., 2012). Furthermore, in beta diversity analysis soil and rhizosphere samples clustered closely together, whereas root and stalk samples separated further apart (**Figure 3A**), indicating stronger compositional divergence from soil-associated communities. The separation along PC1 suggests that the transition from bulk soil to plant-associated compartments (rhizosphere - root - stalk) is accompanied by directional shifts in microbial assemblages, likely reflecting both host filtering and compartment-specific ecological selection.

Plant development also significantly impacts microbial communities with significant shifts in microbiome structure, diversity, and functions as crops progress from early to later developmental (Dadi et al., 2025; Xiong et al., 2021b). Our results show that alpha diversity for soil and rhizosphere compartments remain relatively stable across developmental stages, keeping with prior work showing that soil microbiomes are buffered by stable physical and chemical properties of the soil (Hartmann et al., 2008; Knight et al., 2024). On the other hand, roots and stalk compartments experienced changes in diversity in later growth; this likely reflects the more intimate association between endophytes and plants, and thus a stronger impact of developmental stage. Similar patterns were seen in beta-diversity analysis, although in this case only soil was statistically similar between timepoints (**Figure 3B**). This agrees with prior work showing shifts in the maize rhizosphere community over time but not bulk soil (Walters et al., 2018). Overall, our results show that while progression in plant development significantly restructures the endophytic and plant-associated microbiomes (rhizospheres, roots, and stalk), soil bacterial communities relatively remain stable.

We also assessed the impact of location on alpha and beta diversity of the microbial communities. Interestingly, across different locations, the stalk was the only compartment that exhibited significant differences in the Shannon alpha-diversity metric. Spearman correlation of geographic distance versus UniFrac distances indicate that the relationship between geographic distance and microbial community dissimilarity varies among maize compartments and between developmental stages. Soil, rhizosphere, and root communities showed significant distance-decay relationships at both developmental stages, whereas stalk microbiomes exhibited weak and non-significant spatial structuring at both stages. This suggests that the influence of geographic distance on microbial community assembly changes as the maize plant develops and differs among plant compartments. In a similar study, there was no explanatory value of the spatial distance between the plots for community beta diversity (Grady et al., 2019). This implies that the spatial distance between fields did not explain differences in microbial community composition. That is, plots that were farther apart did not have more different microbiomes than those that were closer together. This contrasts with an analysis of 21 long-term maize fields across China, where community similarity declined with geographic distance (Tan et al., 2020).

### Compartment-specific noise patterns and its implications on microbiome study design

The result from the noise assessment (variability between sample pairs) shows that there are patterns in sampling variability that seem to be specific to each compartment. The rhizosphere microbiome showed the lowest variability, implying this compartment provides the most reproducible measurement for microbiome studies. This possibly reflects differences in colonization dynamics and environmental heterogeneity across these niches. Conversely, the root endosphere exhibits moderate variability, as indicated by a Shannon diversity of approximately 10% and pronounced clustering in beta diversity analyses within pairs. This higher variability (CV 35% for richness) may be due to patchy colonization by endophytes, so that each plant only harbors a subset of available microbes (Hardoim et al., 2015).

Beta diversity showed a spatial clustering of pairs that was consistent across all compartments and the highest beta variability was observed for stalk microbiomes. This is consistent with potentially more stochastic assembly processes and lower overall microbial biomass (as evidenced by the lower rarefaction depth of 500 reads) (Compant et al., 2011). Despite the higher variability observed for stalk, the significant clustering of paired samples (p=0.0069) shows that local environmental filtering, even in sparsely colonized compartments, still exerts detectable effect.

From a practical point of view, these results indicate that single or few samples per location may suffice for rhizosphere and root compartments (but not stalks), though more replicates would of course yield better estimates. This is especially true if richness-based alpha diversity is a focus, since richness-based metrics are quite sensitive to sampling noise (Roswell et al., 2021) and should be interpreted with appropriate statistical controls and replication.

### Bacterial taxa show differential enrichment across timepoints

The abundance of a particular bacterial taxa at different development stages may indicate stage-dependent plant–microbe interactions, nutrient demands, and/or successional dynamics. In our study, there were no significantly differentially abundant ASVs at both timepoints for rhizosphere community when compared to other sample types. This finding contradicts a previous study that shows that both bacterial and fungi taxa were significantly abundant in the rhizosphere than in other compartments (Tebele et al., 2024). Other studies have shown that Proteobacteria have been found to be highly abundant in the rhizosphere and are thought to take advantage of abundant nutrients (Fu et al., 2023; Ling et al., 2022). Similarly, Actinobacteriota are involved in nitrogen cycling, including associations with diazotrophs (nitrogen-fixing bacteria), and are enriched in the rhizosphere compared to bulk soil (Wang et al., 2024b). However, in our study, the genera belonging to these bacterial phyla showed no significant differential abundance. We also observed that the genera significantly abundant at either timepoints (600 and 1400 GDDs) for soil communities belong to the classes Vicinamibacteria, Myxococcia, Alphaproteobateria, and Actinobacteria. Vicinamibacteria has been shown to be drought sensitive or implicated in the formation of dormant cell forms (Huber et al., 2022). In the root, ASVs assigned to genera Massilia, Mucilaginibacter, Sphingomonas, Unclassified Comamonadaceae were significantly abundant at GDD_600 broadly belong to the phyla of Proteobacteria, and Bacteroidota. These three bacteria phyla (Proteobacteria, Actinobacteria, and Bacteroidota) are consistently the most enriched taxa in root and these enrichment pattern have been observed in various crops such as wheat, canola, barley, and soybean (Cordero et al., 2020; Wang et al., 2024a).

### Functional variation of microbial communities

Functional predictions for each community show significant shifts across timepoints. In the rhizosphere, one of the groups of functional predictions focuses on amino acid metabolism (cyanoamino acid metabolism). Maize rhizosphere microbiomes are enriched in gene pathways related to amino acid biosynthesis and metabolism, especially in modern maize lines (Favela et al., 2021; Zhang et al., 2024a), and the same holds for many other plant species (Popchenko et al., 2024; Tashkandi et al., 2022; Wang et al., 2022; Wicaksono et al., 2024; Xia et al., 2021). Amino acids are the main mobilizable source of carbon and nitrogen that support microbial growth in the rhizosphere compared with other chemicals like sugars or organic acids (Moe, 2013). Bacterial amino acids also have roles in both bacterial biofilm formation and disassembly in rhizobacteria (Kolodkin-Gal et al., 2010), an important component of rhizosphere colonization.

In roots, we found significant (albeit not highly abundant) functional genes involving the biosynthesis of secondary metabolites (flavone and flavonol biosynthesis). Many secondary metabolites can shape microbiome composition (Koprivova & Kopriva, 2022) with flavonoids especially prominent (Kudjordjie et al., 2021; Okutani et al., 2020). These metabolites play a part in communication networks and modulate specific functions of the root microbiota (Koprivova & Kopriva, 2022). Flavones and flavonols in root exudates act as signaling molecules, shaping the root microbiome by recruiting beneficial microbes (e.g., diazotrophic bacteria) and modulating biofilm formation, which can enhance biological nitrogen fixation and plant growth (Yan et al., 2022). In maize, root-derived flavones (apigenin and luteolin) mainly enhance the presence of Oxalobacteraceae in the rhizosphere, which positively affects lateral root development and thus boosts maize growth and nitrogen acquisition (Yu et al., 2021). Introducing alleles into maize for advantageous flavones may boost roots’ connections with growth-assisting rhizosphere microbes (Yu et al., 2021). In our study, the stalk microbiome was enriched with functional categories related to metabolism and genetic information processing. These are unfortunately broad categories, and it is difficult to draw conclusions from them, especially with limited information in the literature about bacterial functions in maize stalks.

Together, these predicted functions highlight coordinated microbial roles across maize developmental stages. Future work will be needed to confirm the importance of specific pathways in the different tissue types.

### Community complexity and putative keystone species across sample types

Microbial co-occurrence networks are extensively used to investigate the interconnections within microbial communities because they provide a robust method for investigating the structure, maintenance, and dynamics of these communities, (Faust & Raes, 2012; Fuhrman, 2009; Ren & Gao, 2019). Soil networks consistently exhibited clustering coefficients an order of magnitude higher than their random counterparts, suggesting that the observed network cohesion is not solely an artifact of network size. This was applicable to other sample types as well. Our soil network consistently has higher average degree and clustering coefficients, as well as lower average path length and network diameter, when compared to rhizosphere and root, regardless of the growing degree days. This indicates the complex nature of soil networks compared to these other sample types.

Within community networks, key microorganisms can fill different roles, such as module hubs and connectors (Lupatini et al., 2014). In this study, we identified root as having the most hubs and rhizosphere as having the most connectors respectively at both timepoints. Both classes represent potential keystone species, meaning that if they were removed the modules and networks may also break apart. Such keystone taxa can play essential roles in network structure and, if validated by experimental manipulation, they could be targets for manipulation to improve crop productivity (Olesen et al., 2007). We also saw an increase in the number of module hubs for rhizosphere, root, and stalk as plants developed, while connectors increased in the soil, rhizosphere, and stalk. This implies that as we move from 600 to 1400 GDDs, the number of connectors and module hubs (generalists) increased. More generalists in networks are favorable for maintaining the balance of microbial communities and are important for promoting exchanges of energy, information, and materials among species in networks (Olesen et al., 2007). On the other hand, we saw a reduction in the number of module hubs in soil and connectors in roots. Topologically, network hubs and connectors represent the regulators, mediators or adaptors (Fan et al., 2018). Hence, the loss of connectors and central nodes in modules can result in the breakdown and decline of the entire ecological network (Guimera & Nunes Amaral, 2005). In addition, network size (determined by node and edges) is a poor predictor of hub and connector abundance in these communities. The rhizosphere is not the largest network yet consistently maintains the highest generalist density, and the soil is among the largest networks yet has a relatively low generalist density. Hence, ecological pressures which are specific to each tissue type determine if taxa fill a generalist role.

Within these networks, ASVs from the phyla Proteobacteria and Actinobacteria were generally the dominant keystone taxa. Previous studies have frequently highlighted these clades as keystone taxa in agricultural settings, influencing plant growth and soil nutrient availability (Lupatini et al., 2014; Ma et al., 2016; Xun et al., 2021). We also observed a shift in the role of keystone bacterial taxa over time. For instance, in the soil and rhizosphere, Proteobacteria, Acidobacteriota, and Actinobacteriota served as module hubs at the early timepoint (600 GDDs), while at the later (1400 GDDs) timepoint Actinobacteriota was replaced by Gemmatimonadota. The shift in the role of bacterial taxa was also observed for other sample types and in connector taxa across both timepoints. The role of bacterial taxa in microbial networks changes across different growth stages (Fu et al., 2022; Xiong et al., 2021b; Zhang et al., 2024b). Despite these dynamic changes, some bacterial taxa maintain important roles across multiple crop stages, and this pattern seems to hold for Proteobacteria and Actinobacteriota. Collectively, timepoint drives significant changes in the roles of bacterial taxa within the microbial networks, but some taxa consistently maintain key roles across the two times, supporting both dynamic adaptation and network stability of the microbial network.

### Environmental factors have varying effects on richness and diversity

Of the various environmental factors, we found that humidity consistently impacted soil, rhizosphere, and root microbiome communities. When humidity is high, the soil tends to retain more moisture, which in turn influences species richness and distribution (Jordan et al., 2020). We also found that nitrogen impacts rhizosphere microbial community, agreeing with prior studies that declining soil N availability from plant uptake correlated with corresponding community shifts (Bell et al., 2015).

From our study, we observed that low temperature and wind speed are key climatic variables across root, soil, and rhizosphere bacterial communities’ diversity (**Figure 9**). Windspeed’s impact is not intuitive, though it has previously been found to significantly impact soil microbial diversity (Zhang et al., 2023b); it may serve as a proxy for some other process, like evaporation. Meanwhile, mean annual precipitation and temperature have been shown to serve as indicators of bacterial diversity in maize soil (Tan et al., 2020). We also found that high temperature impacted soil and rhizosphere bacterial community diversity, which agrees with prior work that attributed increase in temperature to changes in bacterial diversity in the rhizosphere (Yu et al., 2019; Zhang et al., 2022).

The soil microbial community is unsurprisingly most impacted by soil properties (Gao et al., 2022). Several physicochemical/soil chemistry factors, including soil clay content, pH, organic carbon, zinc, copper, and salinity, have been identified as influencers of soil microbial diversity (Moffett et al., 2003; Xavier et al., 2019; Zhang et al., 2023b; Zhao et al., 2019). In our study, the key soil chemistry factors that impacted soil bacterial community diversity are K, Cr, Zn, pH, and Pb. In a litchi–peanut intercropping system, higher available K and pH were the main driver of bacterial community variation, explaining compositional differences in RDA at genus and phylum levels (Zhao et al., 2022). In the rhizosphere, Cu emerged as a key determinant of bacterial diversity, consistent with previous findings indicating that both Cu and cadmium (Cd) significantly influence rhizosphere bacterial communities (An et al., 2023). Root microbiome diversity on the other hand had Na as a key factor. Sodium uptake from the soil is a major cause of salinity toxicity (Maathuis & Sanders, 2001), and several studies have shown that bacterial diversity is negatively correlated with soil salinity (Andronov et al., 2012; Guan et al., 2021; Ibekwe et al., 2020; Okie et al., 2015). Most direct studies focus on overall salt (NaCl) rather than sodium alone, however, so it may be that even under nonstress conditions sodium may impact community assembly.

## Conclusion

Our findings in this study provide structural and functional dynamics involving natural microbiomes across different maize growth stages and compartments. Maize compartment, environmental variability, and developmental stage all impacted microbial diversity, reinforcing that these are dynamic, complex systems that are difficult to pin down with a single sample or time point. Although studies like this one provide an important baseline understanding of the makeup and nature of the maize microbiome, identifying which microbes are functionally impacting maize development and how to manipulate them to improve agriculture remains a significant problem. The increasing focus on synthetic communities provides a potentially more tractable window into microbiome manipulation (Parnell et al., 2024; Thoenen et al., 2025), though how well such insights will translate to the much more complex field settings remains to be seen. Future work focusing on functional manipulation of the maize microbial community, especially under commercially relevant conditions, should continue to be a priority for both research and funding agencies.

## Data Availability Statement

All bioinformatics scripts and pipelines are available at https://github.com/Gbemi-bio/Georgia-Microbiome, accessed on 19 November 2026. The raw sequence data are available in the NCBI Sequence Read Archive under accession PRJNA1561722.

