## Supplementary Figure 5 for "Spatiotemporal Dynamics and Environmental Drivers of Maize Microbiome Structure across Diverse Agroecosystems in Georgia, USA"

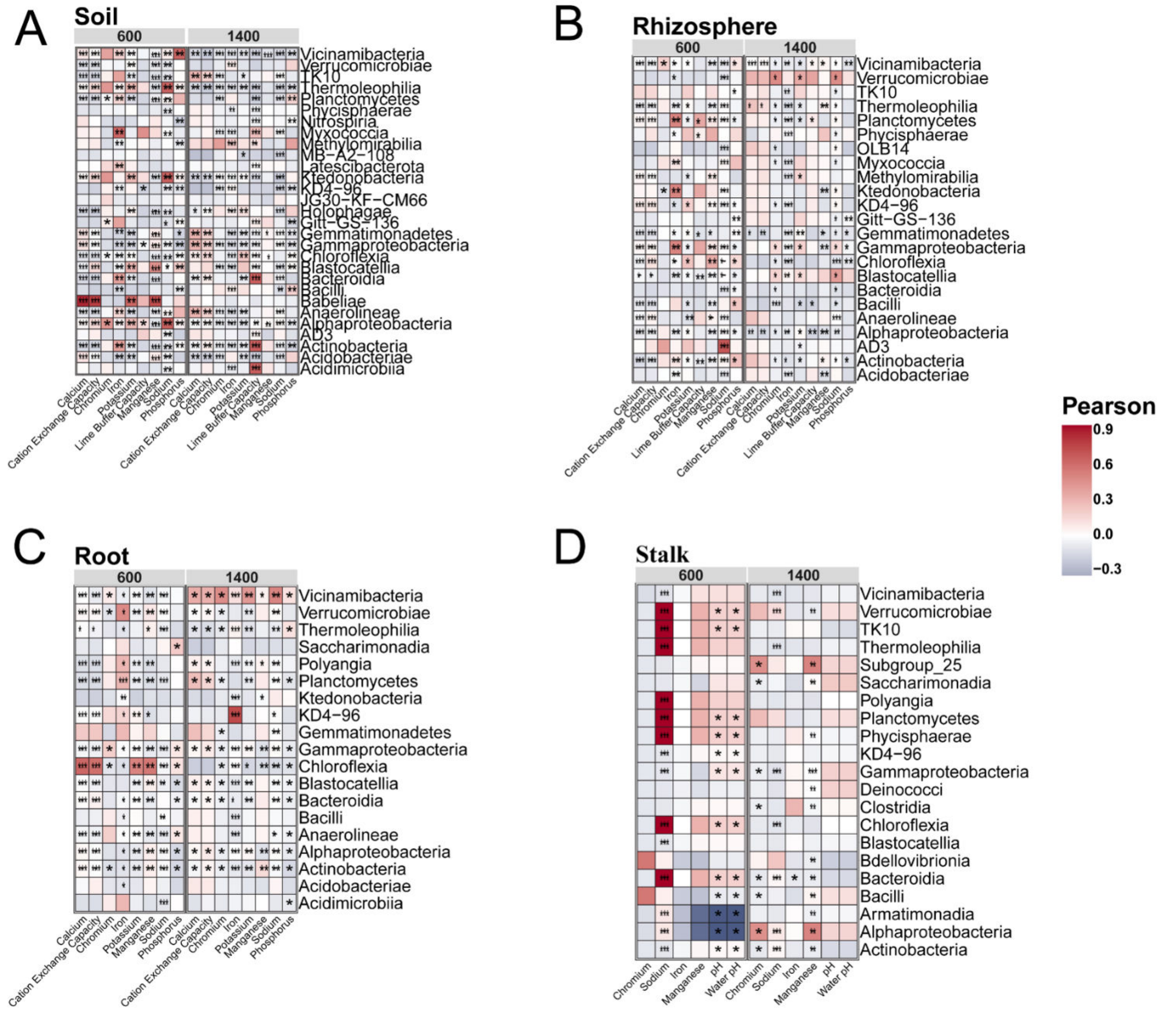
A
Soil
600
1400
Vicinamibacteria
Verrucomicrobiae
TK10
Thermoleophilia
Planctomycetes
Phycisphaerae
Nitrospiria
Myxococcia
Methylomirabilia
MB-A2-108
Latescibacterota
Ktedonobacteria
KD4-96
JG30-KF-CM66
Holophagae
Gitt-GS-136
Gemmatimonadetes
Gammaproteobacteria
Chloroflexia
Blastocatellia
Bacteroidia
Bacilli
Babeliae
Anaerolineae
Alphaproteobacteria
AD3
Actinobacteria
Acidobacteriae
Acidimicrobiia
Calcium
Cation Exchange Capacity
Chromium
Iron
Potassium
Lime Buffer Capacity
Manganese
Sodium
Phosphorus
B
Rhizosphere
600
1400
Vicinamibacteria
Verrucomicrobiae
TK10
Thermoleophilia
Planctomycetes
Phycisphaerae
OLB14
Myxococcia
Methylomirabilia
Ktedonobacteria
KD4-96
Gitt-GS-136
Gemmatimonadetes
Gammaproteobacteria
Chloroflexia
Blastocatellia
Bacteroidia
Bacilli
Anaerolineae
Alphaproteobacteria
AD3
Actinobacteria
Acidobacteriae
Pearson
0.9
0.6
0.3
0.0
−0.3
C
Root
600
1400
Vicinamibacteria
Verrucomicrobiae
Thermoleophilia
Saccharimonadia
Polyangia
Planctomycetes
Ktedonobacteria
KD4-96
Gemmatimonadetes
Gammaproteobacteria
Chloroflexia
Blastocatellia
Bacteroidia
Bacilli
Anaerolineae
Alphaproteobacteria
Actinobacteria
Acidobacteriae
Acidimicrobiia
D
Stalk
600
1400
Vicinamibacteria
Verrucomicrobiae
TK10
Thermoleophilia
Subgroup_25
Saccharimonadia
Polyangia
Planctomycetes
Phycisphaerae
KD4-96
Gammaproteobacteria
Deinococci
Clostridia
Chloroflexia
Blastocatellia
Bdellovibrionia
Bacteroidia
Bacilli
Armatimonadia
Alphaproteobacteria
Actinobacteria
Chromium
Sodium
Iron
Manganese
pH
Water pH
