## Supplementary Figure 1 for "Spatiotemporal Dynamics and Environmental Drivers of Maize Microbiome Structure across Diverse Agroecosystems in Georgia, USA"

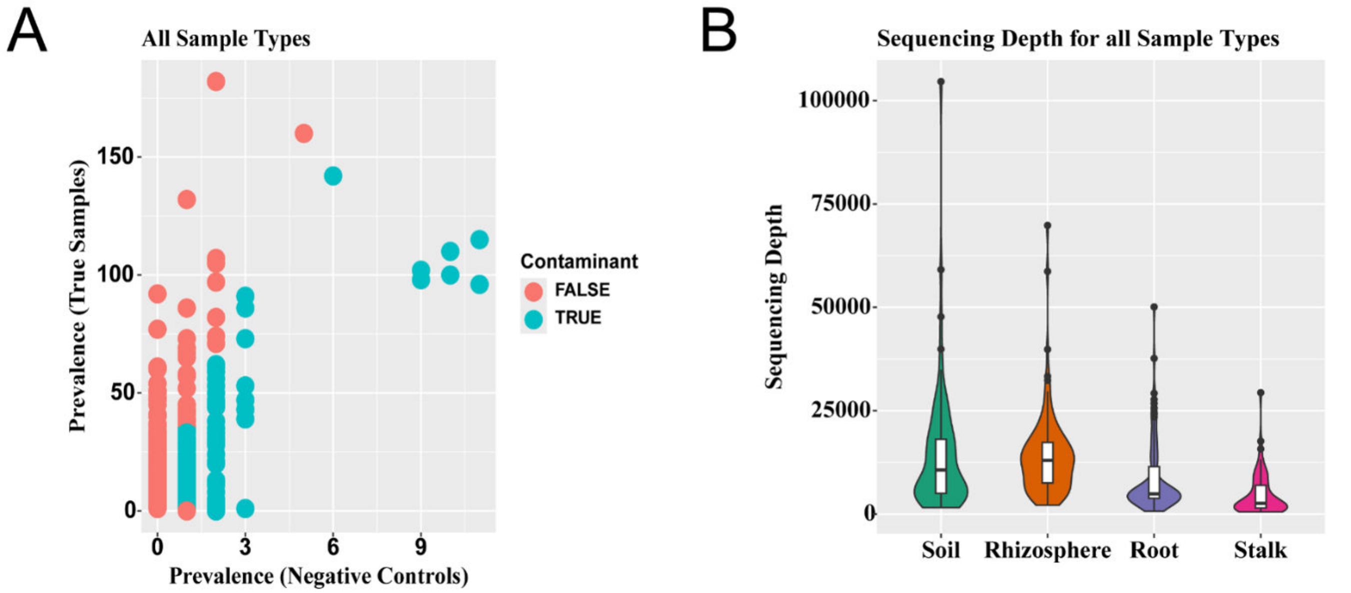
A
All Sample Types
Prevalence (True Samples)
150
100
50
0
0
3
6
9
Prevalence (Negative Controls)
Contaminant
FALSE
TRUE
B
Sequencing Depth for all Sample Types
Sequencing Depth
100000
75000
50000
25000
0
Soil
Rhizosphere
Root
Stalk
