## Supplementary Figure 2 for "Spatiotemporal Dynamics and Environmental Drivers of Maize Microbiome Structure across Diverse Agroecosystems in Georgia, USA"

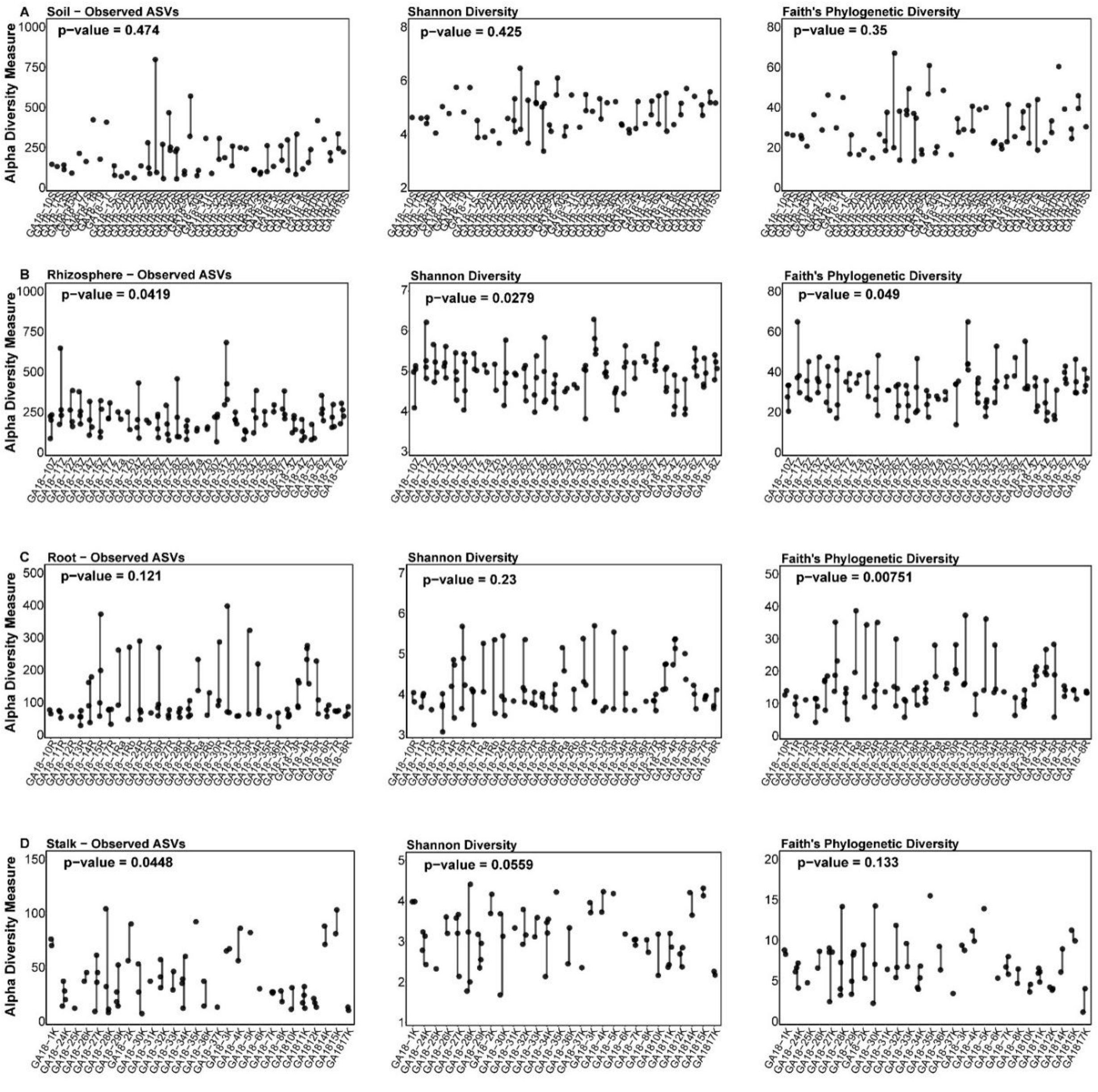
A
Soil − Observed ASVs
p−value = 0.474
Shannon Diversity
p−value = 0.425
Faith's Phylogenetic Diversity
p−value = 0.35
B
Rhizosphere − Observed ASVs
p−value = 0.0419
Shannon Diversity
p−value = 0.0279
Faith's Phylogenetic Diversity
p−value = 0.049
C
Root − Observed ASVs
p−value = 0.121
Shannon Diversity
p−value = 0.23
Faith's Phylogenetic Diversity
p−value = 0.00751
D
Stalk − Observed ASVs
p−value = 0.0448
Shannon Diversity
p−value = 0.0559
Faith's Phylogenetic Diversity
p−value = 0.133
Alpha Diversity Measure
0 250 500 750 1000
2 4 6 8
0 20 40 60 80
3 4 5 6 7
0 100 200 300 400 500
0 10 20 30 40 50
0 50 100 150
1 2 3 4 5
0 5 10 15 20
