## Supplementary Figure 3 for "Spatiotemporal Dynamics and Environmental Drivers of Maize Microbiome Structure across Diverse Agroecosystems in Georgia, USA"

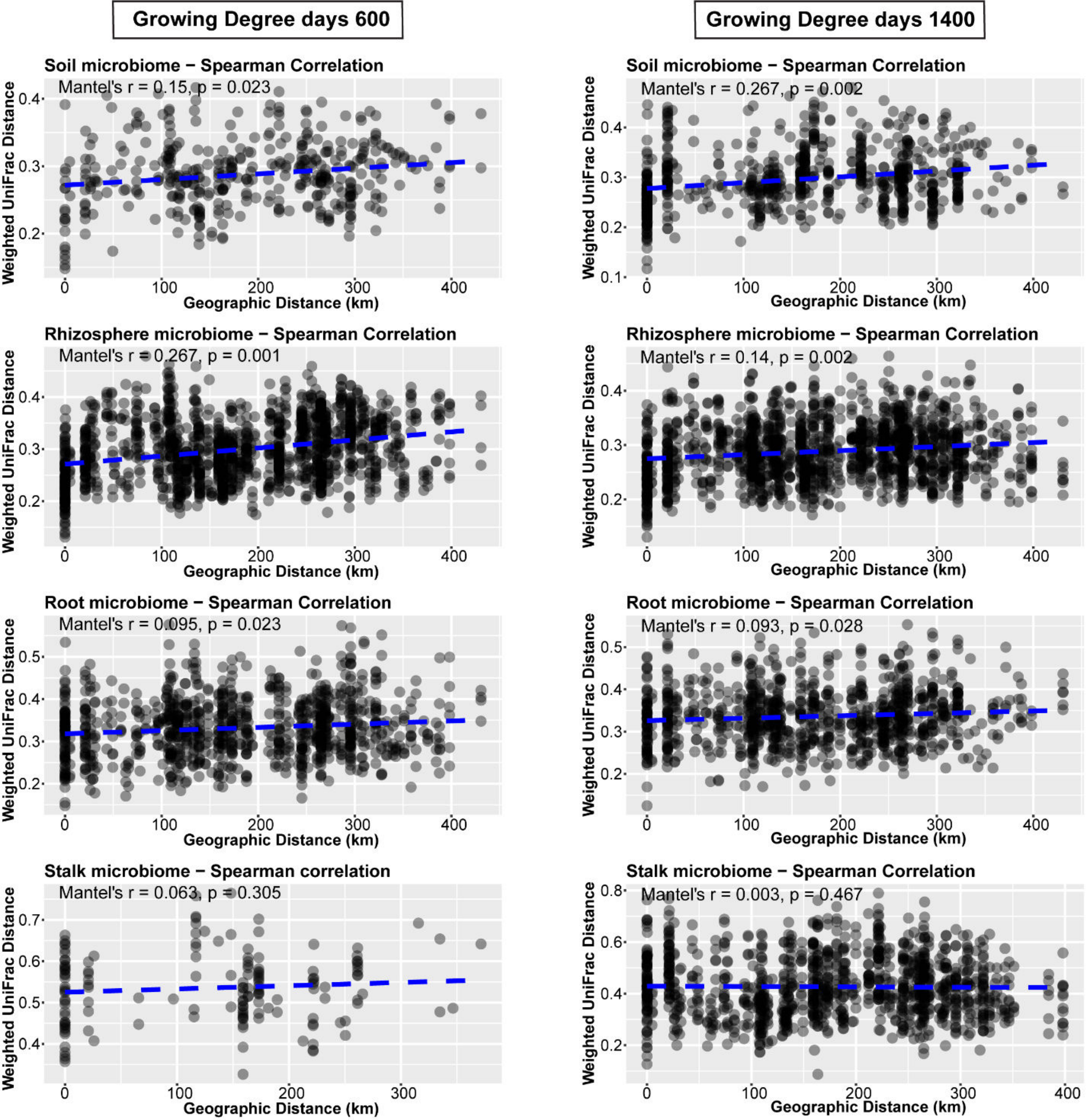

Growing Degree days 600
Growing Degree days 1400
Soil microbiome − Spearman Correlation
Mantel's r = 0.15, p = 0.023
Soil microbiome − Spearman Correlation
Mantel's r = 0.267, p = 0.002
Rhizosphere microbiome − Spearman Correlation
Mantel's r = 0.267, p = 0.001
Rhizosphere microbiome − Spearman Correlation
Mantel's r = 0.14, p = 0.002
Root microbiome − Spearman Correlation
Mantel's r = 0.095, p = 0.023
Root microbiome − Spearman Correlation
Mantel's r = 0.093, p = 0.028
Stalk microbiome − Spearman correlation
Mantel's r = 0.063, p = 0.305
Stalk microbiome − Spearman Correlation
Mantel's r = 0.003, p = 0.467
Weighted UniFrac Distance
Geographic Distance (km)
