## Supplementary Figure 4 for "Spatiotemporal Dynamics and Environmental Drivers of Maize Microbiome Structure across Diverse Agroecosystems in Georgia, USA"

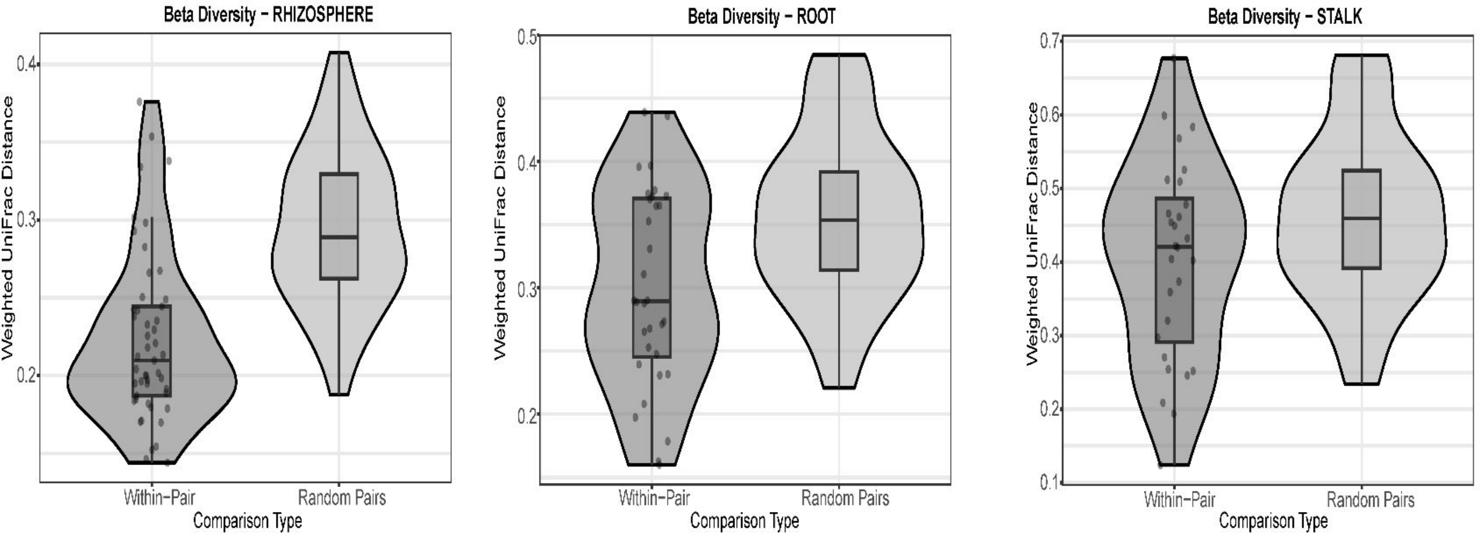
Beta Diversity – RHIZOSPHERE
Weighted UniFrac Distance
0.4
0.3
0.2
Within–Pair
Random Pairs
Comparison Type
Beta Diversity – ROOT
0.5
0.4
0.3
0.2
Beta Diversity – STALK
0.7
0.6
0.5
0.1
